# Somatosensory high frequency oscillations across the human central nervous system

**DOI:** 10.1101/2024.11.16.622608

**Authors:** Emma Bailey, Birgit Nierula, Tilman Stephani, Gunnar Waterstraat, Gabriel Curio, Vadim Nikulin, Falk Eippert

## Abstract

Information transmission in the central nervous system (CNS) relies on the propagation of action potentials (neuronal spikes), traditionally thought to be inaccessible with electroencephalography (EEG). Converging evidence from human and animal recordings however demonstrates that high frequency oscillations (HFOs) in EEG recordings represent neuronal population spiking, complementing standard low-frequency EEG recordings in cortex. Here, we developed an approach to simultaneously record HFOs to somatosensory stimulation across the entire human CNS noninvasively: from spinal over subcortical to cortical stages. Our approach provides a detailed temporal and spectral characterization of HFOs at each CNS level, delineating how they evolve across the somatosensory processing hierarchy. Critically, we show that low and high frequency electrophysiological responses represent at least partly independent processing channels across the CNS. We thus demonstrate the feasibility of studying the full spectrum of neuronal information processing across the CNS, offering a multi-level window into human neurophysiology in health and disease.

## Introduction

Methods for non-invasive and time-resolved recordings of brain function are important tools in human neuroscience and generally fall into two classes: those using indirect read-outs of neuronal function via hemodynamic changes (e.g. functional magnetic resonance imaging) and those that record direct manifestations of neuronal processes (e.g. electroencephalography [EEG] or magnetoencephalography [MEG]). Despite their excellent temporal resolution, the latter largely focus on neuronal signals with frequencies below ∼100Hz – thought to reflect mostly postsynaptic potentials (Baillet, 2017; Biasiucci et al., 2019) – although much faster neuronal signals are ubiquitous throughout the brain.

For example, in the somatosensory domain, high frequency oscillations (HFOs) co-occur with low-frequency somatosensory evoked potentials (LF-SEPs) after peripheral nerve stimulation (Abbruzzese et al., 1978; Cracco and Cracco, 1976) and have been studied with human EEG (Eisen et al., 1984; Yamada et al., 1988) and MEG (Curio et al., 1994; Hashimoto et al., 1996). Importantly, animal EEG/MEG recordings linked HFOs at ∼600Hz to spiking activity in somatosensory cortex (Baker et al., 2003; Ikeda et al., 2002; Telenczuk et al., 2011), which is of high interest considering the critical role spikes play in neuronal information transfer. Since recording cortical spike activity in humans is possible only invasively in the context of neurosurgical procedures, HFO recordings provide a unique non-invasive window on cortical population spikes, alongside the concurrently acquired low-frequency signals of post-synaptic origin.

Although the precise cellular origins of cortical HFOs are still debated (Coppola, 2004; Gobbelé et al., 2007; Hu et al., 2023; Inoue et al., 2004, 2001; Restuccia et al., 2003), they are at least partly independent of their low-frequency counterparts, with differing effects of stimulus repetition, movement, sleep and anaesthesia (Hashimoto et al., 1996; Inoue et al., 2002; Klostermann et al., 2000; Restuccia et al., 2011). Furthermore, high-frequency neural signals are not limited to electrical somatosensory nerve stimulation, but are also observed with tactile (Baker et al., 2003; Barth, 2003) and visual input (Bishop and Clare, 1953; Funke and Kerscher, 2000) as well as motor cortex stimulation (Beck et al., 2023; Lazzaro et al., 1998). Additionally, HFOs appear to be a processing mode of the entire central nervous system (CNS): non-invasive support comes from dipole source analysis revealing generators in the brainstem, thalamus and cortex (Gobbelé et al., 2007; Halboni et al., 2000), nasopharyngeal recordings showing dorsal column pathway responses (Restuccia et al., 2004), and neck recordings demonstrating cervical spinal cord responses (Chander et al., 2022; Stenner et al., 2025); invasive data provide further support (Berić et al., 1986; Hanajima et al., 2004; Insola et al., 2010, 2008; Jeanmonod et al., 1991; Stenner et al., 2025; Urasaki et al., 2006).

Yet, there is a complete lack of systemic approaches that would allow for concurrently recording and characterizing HFOs across the entire CNS. One reason for this is the difficulty in recording them: HFOs are over 10 times smaller than LF-SEPs (Nakano and Hashimoto, 1999) and their recording benefits from specialised low-noise recording setups (Waterstraat et al., 2021, 2016), but these are currently limited to cortical recordings. Entire-CNS HFO recordings may however be crucial for uncovering systematic changes in the spectral and temporal aspects of neuronal information propagation, also with respect to HFO contributions to pathology (e.g. in epilepsy, focal dystonia and Parkinson’s disease (Cimatti et al., 2006; Yang et al., 2014; Zijlmans et al., 2012)).

Here, we therefore leveraged advances in non-invasive multi-channel electrophysiology (Bailey et al., 2024; Nierula et al., 2024; Waterstraat et al., 2015) to simultaneously record somatosensory HFOs from spinal, subcortical and cortical levels, i.e. across the entire CNS processing hierarchy. We demonstrate the robustness of our approach (via replication across two studies) as well as the generalizability of its insights (by using both upper and lower limb stimulation). Considering the link between HFOs and spiking (Baker et al., 2003; Ikeda et al., 2002; Telenczuk et al., 2011), this approach provides a new window into neuronal signalling across the entire CNS, concurrently offering insights into fast (spiking) and slow (postsynaptic) neuronal processes.

## Results

Here, we utilized two independent datasets (Dataset 1: N=36; Dataset 2: N=24) of multichannel surface recordings across the spinal cord and brain in response to unilateral electrical stimulation of the left median and tibial nerves (reflecting upper and lower limb responses, respectively). To improve sensitivity, we leveraged the multivariate nature of these data by using a spatial filtering approach based on canonical correlation analysis (CCA).

### Single participant analyses

Considering that HFOs have not yet been recorded simultaneously across the CNS, we first assessed the ability of our approach to detect HFO bursts at the level of single participants. Representative individual timecourses (Dataset 1: Figure 1; Dataset 2: Figure S1) demonstrated a distinct HFO burst at each level of the CNS (spinal, subcortical and cortical) with a latency similar to the LF-SEP peak, suggesting sufficient data quality.

**Figure 1.**
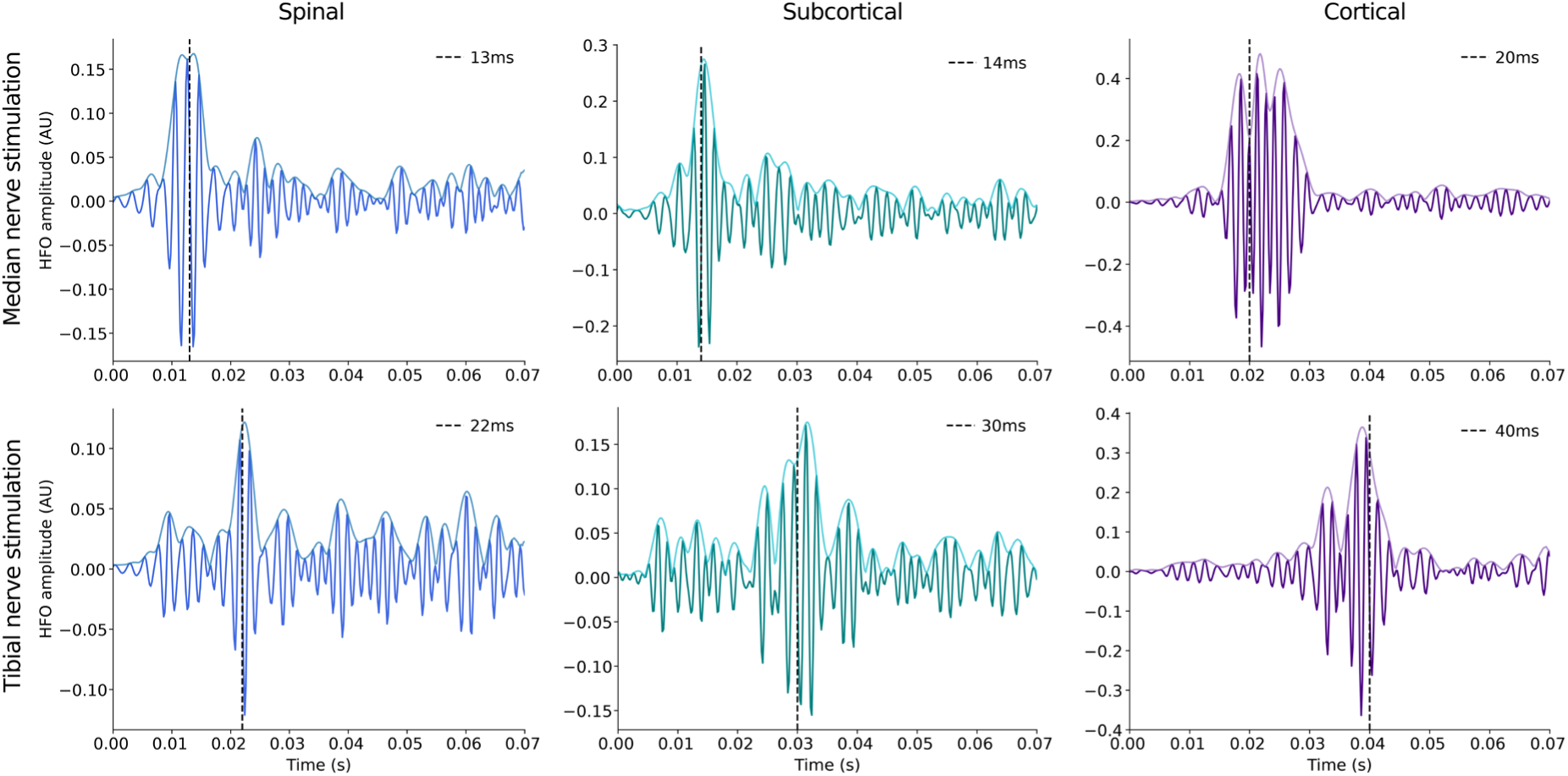
Individual HFO bursts across the CNS. Single participant HFO timecourses after CCA (thick lines) and their amplitude envelopes (thin lines) in response to median nerve stimulation (participant 3: top) and tibial nerve stimulation (participant 31: bottom) across spinal (left), subcortical (middle) and cortical (right) recordings. For each participant, condition and CNS level, the optimal CCA spatial filter was applied to the sensor-space data and plotted in the time domain as shown here. For each panel, the canonical latency of the LF-SEP is displayed as a dashed vertical line. All data from Dataset 1; a corresponding depiction of data from Dataset 2 can be found in Figure S1.

This observation was quantitatively supported by the overall proportion of participants in which we were able to identify a somatosensory HFO CCA component (Table 1a): for median nerve stimulation, we were able to identify CCA components at every CNS level in more than 90% of participants and for tibial nerve stimulation a similar proportion emerged at cortical and subcortical levels, with lower proportions at the spinal level (52% on average). The advantage of spatial filtering is clearly evident when comparing these numbers to those resulting from aiming to detect the HFO burst in sensor space (i.e. prior to the application of the optimal spatial filter; Table 1b): this is almost impossible for tibial nerve stimulation (at most 19% in cortex) and suboptimal even for median nerve stimulation (at most 75% in cortex), with the worst performance at the spinal level. Together, these data demonstrate that spinal, subcortical and cortical HFOs are a robust phenomenon, detectable non-invasively in a large majority of healthy participants, provided that adequate analysis techniques are employed.

**Table 1.**
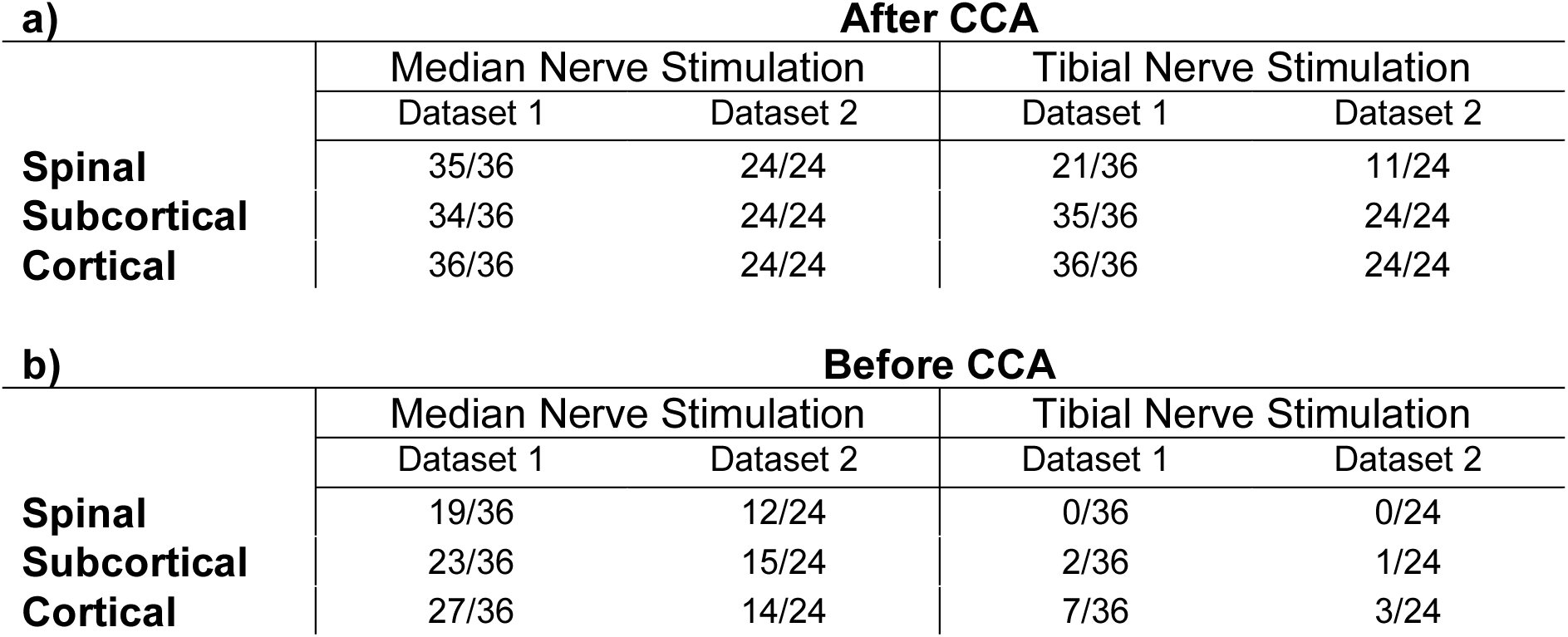
Proportion of participants with detectable HFOs before and after CCA. Proportion of participants in which selection criteria were satisfied a) after CCA (i.e. for at least one CCA component) or b) before CCA (i.e. within the channel of interest in sensor space prior to the application of the CCA spatial filters). The rows depict the CNS level and the columns the different datasets. Note that selection was automated and thus not influenced by possible experimenter bias.

### Group level analyses

#### Timecourses and topographies

We next aimed for a spatiotemporal HFO characterization across the CNS, first assessing group-level responses in the time domain, for which it was necessary to average the HFO amplitude envelopes across participants. Both for median and tibial nerve stimulation we observed clear group-level HFO envelope timecourses (Figure 2), with peaks at latencies that are in accordance with the CNS level: median nerve stimulation HFO peaks at 10.6ms (spinal), 15.2ms (subcortical) and 17.8ms (cortical), and tibial nerve stimulation HFO peaks at 22.4ms (spinal), 31.0ms (subcortical) and 40.4ms (cortical); Figure S2 and Supplementary Results show replication in Dataset 2. For spinal data, it is interesting to note that although the spinal HFO peak occurs before the canonical low frequency response to median nerve stimulation at 13ms, the envelope timecourse is clearly above noise level until after this timepoint. For subcortical data, it is important to point out that the observed peak – with a latency between that of spinal and cortical responses – is not due to the selected CCA training window (i.e. possible overfitting), but can also be observed in sensor-space data (Figure S3).

**Figure 2.**
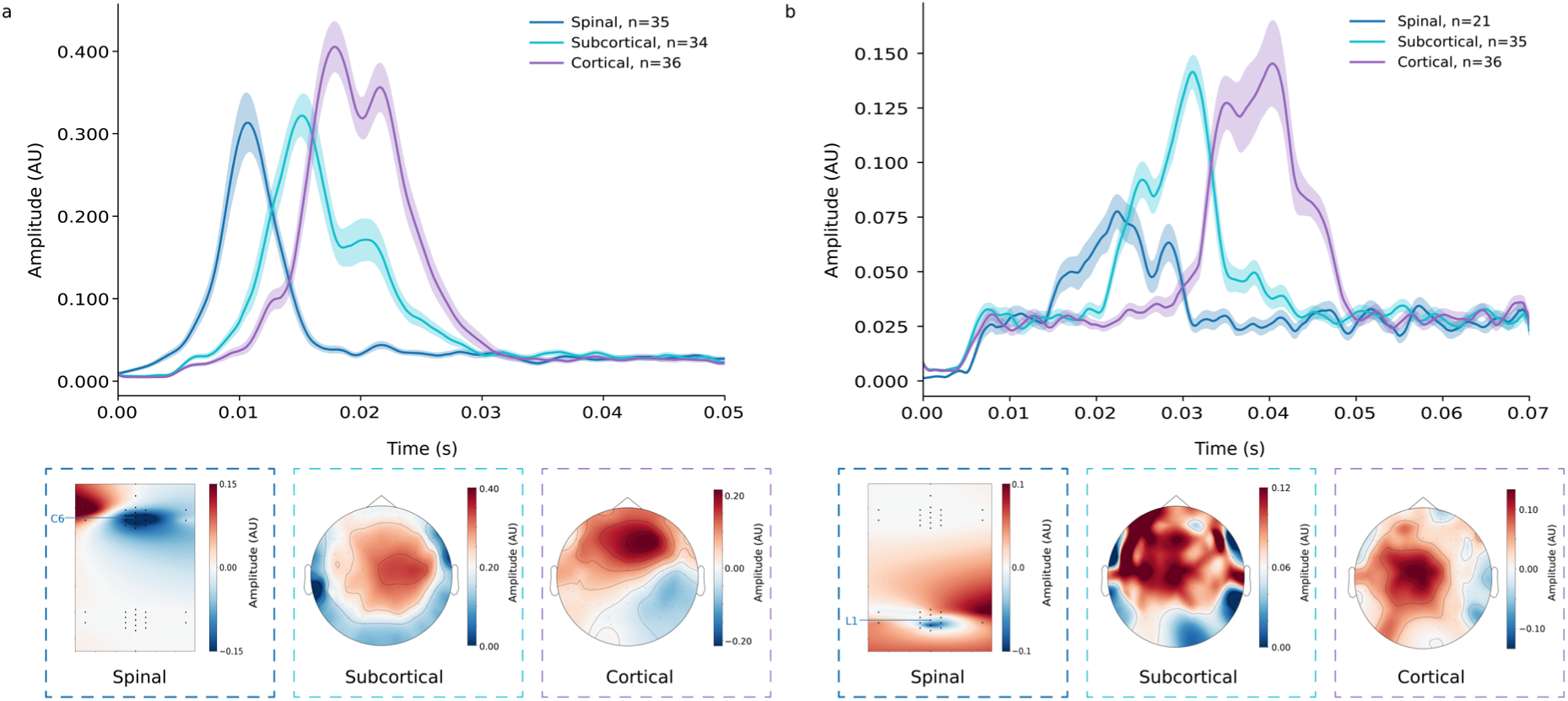
Group-level HFO timecourses and topographies across the CNS. Depicted are group-averaged HFO amplitude envelopes and the associated spatial activation patterns for spinal (black dots representing channel locations), subcortical and cortical responses to a) median nerve stimulation and b) tibial nerve stimulation after the application of CCA. The error bands reflect the SEM across participants and the number of eligible participants entering each average is given in the legend. Please also note that different scales are applied to each plot. All data from Dataset 1; a corresponding depiction of the high frequency responses from Dataset 2 can be found in Figure S2.

The group-level spatial topographies (Dataset 1: Figure 2; Dataset 2: Figure S2) aligned with the expected spatial pattern based on the stimulated nerve and central nervous system level and were highly similar to low-frequency topographies (Figure S4). Of note, spinal topographies exhibited an additional feature that is not observed in low-frequency data (but replicable across datasets): the cervical response to median nerve stimulation showed an additional lateralized response peak that is ipsilateral to the side of stimulation and could point towards a brachial plexus contribution.

Finally, as CCA is an optimisation algorithm which will enhance signals in the chosen time period even in the absence of true stimulus-evoked neural activity, we performed an additional validation analysis to determine whether CCA extracted genuine stimulus-evoked HFOs. Here, we observed that detection of HFOs was consistently more reliable in stimulation-based data compared to resting-state data obtained from the same participants (where no stimulation-evoked HFOs should exist; Supplementary Results and Table S1).

#### Time-frequency representation

Next, we turned to a group-level spectral characterization of HFOs, which is established at the cortical level, but so far elusive across the CNS. Time-frequency representations (TFRs) were obtained for the evoked response to both median and tibial nerve stimulation at each level of the CNS after applying the optimal spatial filter (Dataset 1: Figure 3; Dataset 2: Figure S5). TFRs for both median and tibial nerve stimulation showed areas of increased power relative to baseline in the frequency band of interest (400-800Hz), and at time points relevant to spinal, subcortical and cortical responses. Importantly, although broadband data are represented (i.e. no spectral filtering having taken place), the high-frequency peaks are clearly distinct from low-frequency activity, with the exception of spinal responses to tibial nerve stimulation where LF and HF bleed into another in Dataset 1 (Figure 3), though not in Dataset 2 (Figure S5). One noteworthy feature for median nerve stimulation is the frequency of HFOs appearing to increase along the processing hierarchy from spinal over subcortical to cortical responses, which we next assessed quantitatively.

**Figure 3.**
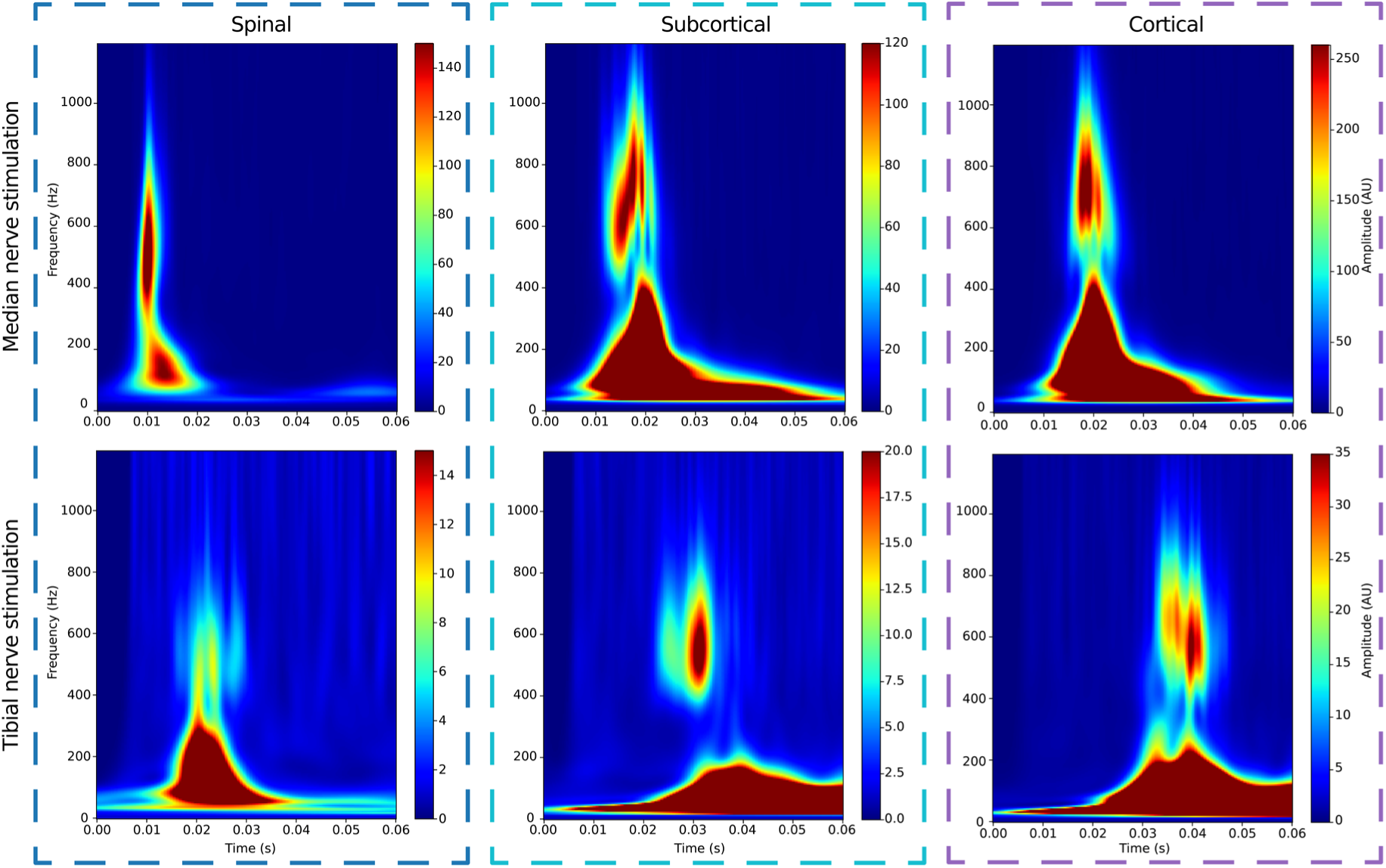
Evoked power across the CNS. Group-average time-frequency representations of the evoked response to median nerve (top) and tibial nerve (bottom) stimulation, including spinal (left), subcortical (middle) and cortical (right) responses. Please note the different scales applied to each time-frequency plot. All data from Dataset 1; a corresponding depiction of data from Dataset 2 can be found in Figure S5.

#### Burst frequency

We therefore calculated the HFO burst frequency for each participant and CNS level (for group-level results, see Table 2). Descriptively, there is an increase in HFO burst frequency as the central nervous system is ascended towards the cortical level – this pattern held across both data-sets and both stimulated nerves with the exception of spinal responses to tibial nerve stimulation (where variability across participants is much stronger, see Table 2). Statistical testing via repeated measures ANOVAs and follow-up t-tests largely supported these descriptive results and revealed significant differences between the mean burst frequency at different CNS levels in response to median nerve stimulation in both datasets, with no significant differences found after tibial nerve stimulation (see Supplementary Results).

**Table 2.** Group-average burst frequency in Hertz across the central nervous system, expressed as mean ± SEM across participants.

|  | **Median Nerve Stimulation** |  | **Tibial Nerve Stimulation** |  |
| --- | --- | --- | --- | --- |
|  | Dataset 1 | Dataset 2 | Dataset 1 | Dataset 2 |
| **Spinal** | 528.3 ± 4.7 | 524.5 ± 5.3 | 562.7 ± 9.2 | 580.2 ± 10.3 |
| **Subcortical** | 590.8 ± 7.1 | 576.8 ± 7.6 | 552.6 ± 6.8 | 557.1 ± 7.4 |
| **Cortical** | 652.3 ± 6.8 | 646.4 ± 6.1 | 579.6 ± 7.0 | 580.2 ± 7.9 |

#### Wavelet peak count

Since the number of peaks within a cortical HFO wavelet has been related to the number of peaks in single-unit post stimulus time histograms of cortical neurons (Baker et al., 2003; Telenczuk et al., 2011), we next investigated whether the number of peaks within an HFO wavelet would differ across the CNS levels (Figure 4 and Figure S6; note that simulations excluded the possibility of filter ringing influencing these results, as shown in Supplementary Results). As an example, for median nerve stimulation we observed a group-average of 4-8 peaks cortically, while only 2-6 peaks were evident at the spinal level. Visual inspection of median nerve stimulation results showed an increase in the number of peaks from spinal over subcortical to cortical levels, while there was no clear trend for responses to tibial nerve stimulation. Both of these observations were supported statistically, and consistent across both datasets (Supplementary Results).

**Figure 4.**
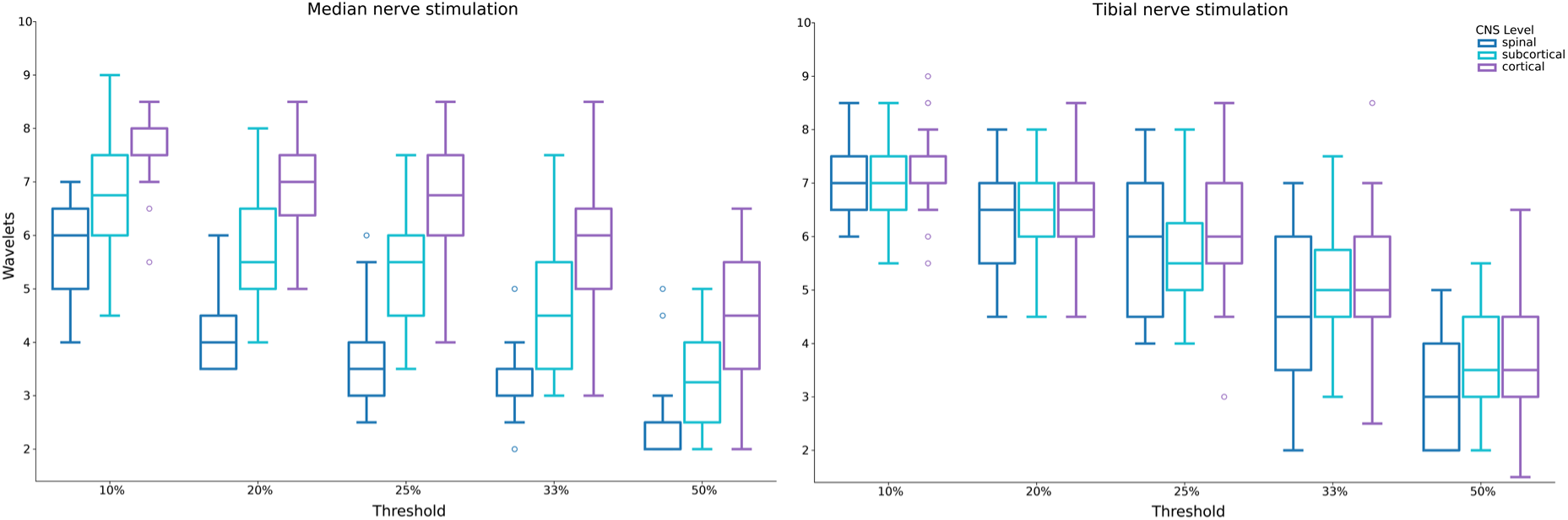
Wavelet peak count across the CNS. Depicted are the group-average number of wavelet peak bursts at each level of the CNS, dependent on different thresholds used for peak-counting (i.e. relation of peak/trough size to maximum peak/trough, respectively). All data from Dataset 1; a corresponding depiction of data from Dataset 2 can be found in Figure S6.

#### Sensory nerve stimulation

We next asked whether our approach would also allow for detecting HFOs in a low SNR scenario, using the sensory nerve stimulation conditions of Dataset 2 (finger and toe stimulation), where a much lower number of nerve fibres is stimulated. Automated component selection revealed that there were sufficient participants for statistical analysis of cortical (18/24) and subcortical responses (7/24) to finger stimulation; but neither spinal responses to finger stimulation (2/24) nor any response to toe stimulation (0/24 for all three levels) had enough participants with automatically detected HFOs.

While there was large inter-individual variability in the strength of the sensory-nerve HFOs (Figure 5), cluster-based permutation testing revealed that it is possible to robustly detect group-level cortical responses after single (finger1, finger2) and double digit (finger12) stimulation, as well as subcortical responses after double digit stimulation (Supplementary Results). Thus, while such sensory-nerve HFOs are more difficult to detect, they can still be observed with our approach, and meaningful aspects can be revealed (e.g. earlier clusters for subcortical than cortical sensory nerve responses).

**Figure 5.**
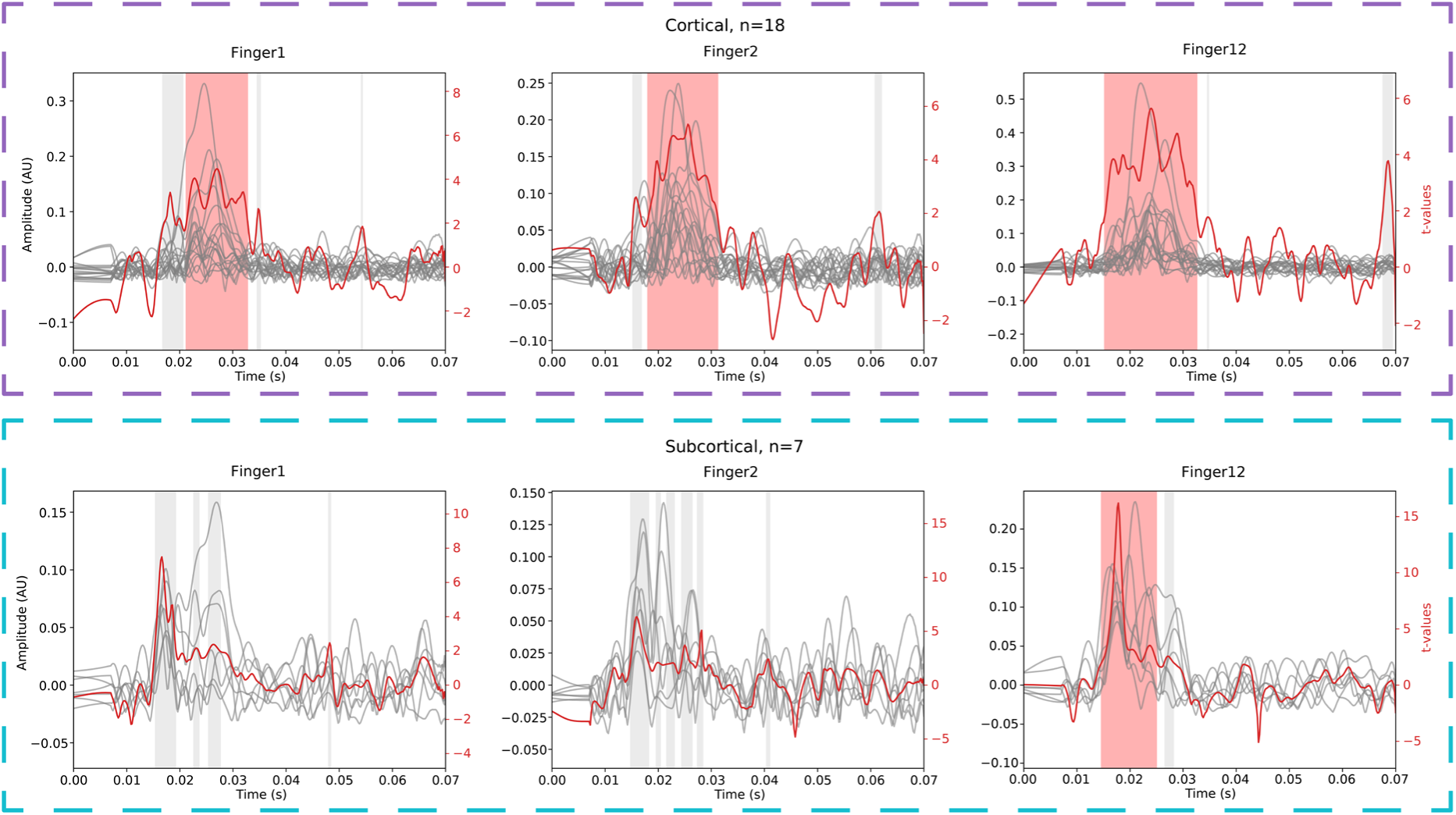
Sensory nerve HFOs. Gray lines depict participants’ individual cortical (top) and subcortical (bottom) amplitude envelopes after single (finger1, finger2) or double (finger12) digit stimulation. The red line depicts the temporally resolved t-values from the cluster-based permutation tests (scale on right side of plot), with the shaded areas indicating non-significant (grey) and significant (red) clusters at p < 0.05. Please note the different y-axis scaling used in each panel.

#### Relationship between low-frequency and high-frequency responses

In a final set of analyses, we aimed to probe in two complementary ways – assessing between-participant and within-participant variation – whether low-frequency and high-frequency responses contain redundant information or function as rather independent processing channels.

### Correlation between average LF-SEP and HFO peak amplitude

First, we investigated whether there would be a significant correlation between average LF-SEP and HFO peak amplitudes across participants, which would speak for these signals containing redundant information. While this was the case in many instances (Supplementary Results), when accounting for possibly confounding effects of SNR (by considering the partial correlation controlling for LF-SEP SNR, HFO SNR or both), only one out of the eight investigated correlations across both datasets showed significant results (Supplementary Results). Thus, the vast majority of analyses showed these signals contain non-redundant information.

### Comparing LF-SEP and HFO timecourses when considering the strongest versus the weakest trials

Second, we investigated this question at the level of single-trials by selecting the top and bottom 10% of trials based on HFO SNR for each participant and then assessing differences in the evoked response for both LF-SEP and HFO; in case of redundant information, single-trial LF-SEP and HFO amplitudes should be similarly high or low. Using this approach, it was evident that while there was a large difference between the HFO amplitude envelope produced using the strongest versus the weakest 200 trials, there was no perceptible difference between the LF-SEPs when using the same ranking of trials (Figure 6; note that the HFO difference is to be expected, considering that selection and testing are not independent here; the important finding is the absence of differences in the LF-SEP). This pattern was statistically confirmed via cluster-based permutation tests at both the cortical and spinal level (Supplementary Results). Further, in a complementary analysis based on ranking according to the LF-SEP SNR, the opposite effect was observed whereby there was a statistically significant difference between the strongest and weakest LF-SEPs, but not the corresponding HFOs (Figure S7). Both types of results were furthermore replicated in Dataset 2 (Figure S8 and S9, as well as Supplementary Results). Together, these results suggest that LF and HF responses represent at least partly independent information channels.

**Figure 6.**
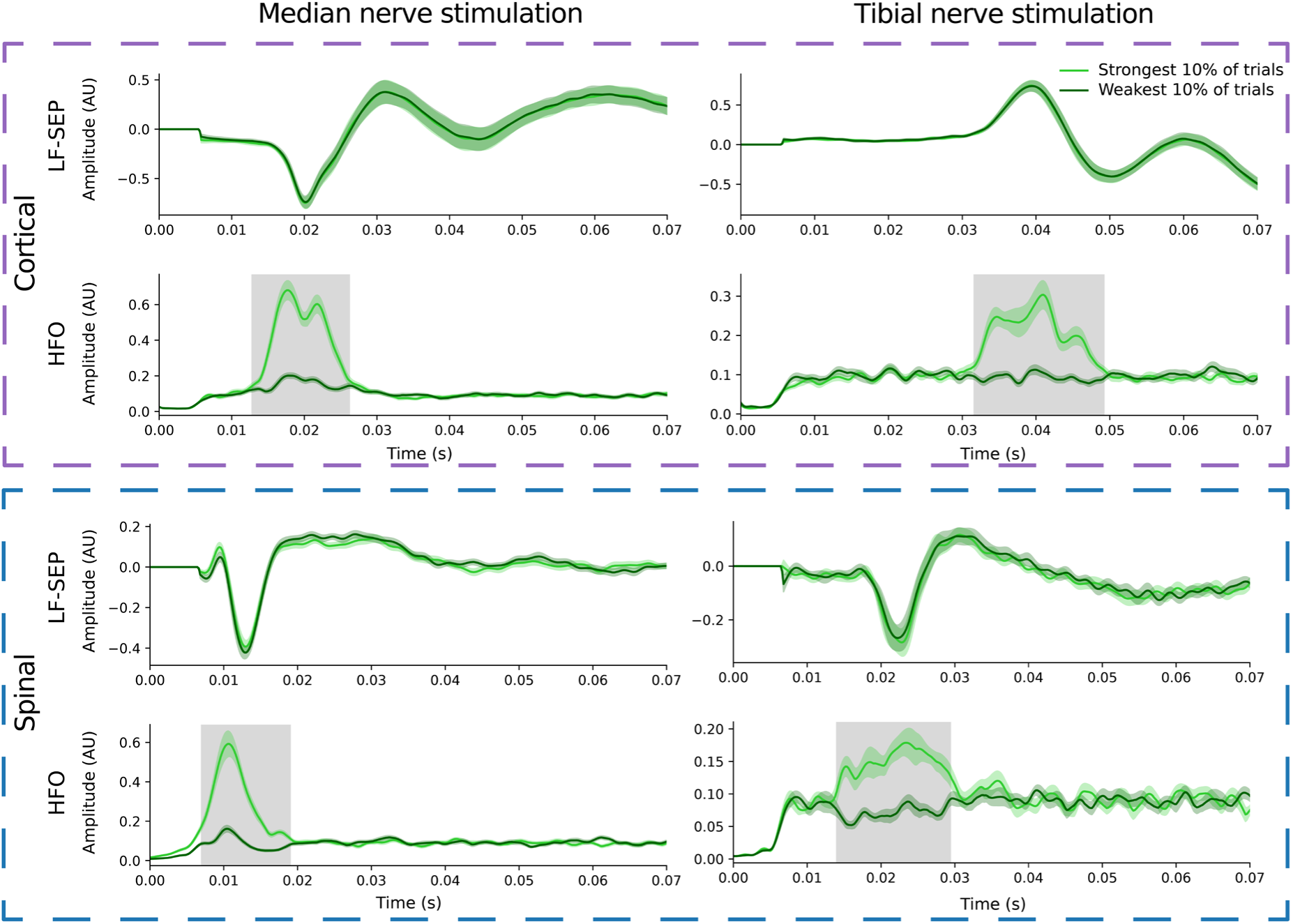
Difference between low-frequency and high-frequency signals. Depicted are the group-average low-frequency somatosensory evoked potentials (LF-SEP) and group-average high-frequency amplitude envelopes (HFO) when considering either the strongest (light green) or weakest (dark green) 10% of trials (as ranked according to the HFO SNR). The upper two panels show cortical data after median (left; n=36) and tibial (right; n=36) nerve stimulation, whereas the lower two panels show spinal data (median nerve: n=35; tibial nerve: n=20). The grey areas indicate significant clusters at p < 0.05 from a cluster-based permutation test. Please note the different y-axis scales employed in each plot. All data from Dataset 1; a corresponding depiction of data from Dataset 2 can be found in Figure S8.

## Discussion

In this study, we systematically assessed somatosensory high frequency oscillations (HFOs) to peripheral nerve stimulation across the central nervous system (CNS). While the use of two independent datasets allowed for an internal replication of the obtained results and thus ensured their robustness, the stimulation of both the median nerve (upper limb) and the tibial nerve (lower limb) allowed us to assess the generalizability of our findings. Combining EEG recordings with spinal recordings (via dense multi-channel electrode grids), we observed that it is possible to non-invasively record HFOs in a frequency range of 400-800 Hz – previously linked to population spikes in cortex – from cortical, subcortical and spinal levels, providing a novel and comprehensive window into human neurophysiology.

### Feasibility of non-invasive HFO recordings across the CNS

Although HFOs were discovered almost five decades ago (Abbruzzese et al., 1978; Cracco and Cracco, 1976), it only became apparent in the last two decades that they could provide a novel window into neuronal ensemble activity when their relation to population spikes in cortex was discovered (Baker et al., 2003; Ikeda et al., 2002; Telenczuk et al., 2011). Recent advances in spatial filtering techniques and low-noise data acquisition systems – both developed to address the low SNR of HFOs – have since rekindled interest in these non-invasive macroscopic signals in human neuroscience (Chander et al., 2022; Stenner et al., 2025; Stephani et al., 2024; Waterstraat et al., 2021).

Here, we demonstrate that it is possible to obtain somatosensory HFOs simultaneously across the entire CNS processing hierarchy – from spinal over subcortical to cortical processing sites – using readily available EEG equipment. This represents a critical step forward, as previous invasive (Hanajima et al., 2004; Insola et al., 2010, 2008; Stenner et al., 2025; Urasaki et al., 2006) and non-invasive HFO studies (Chander et al., 2022; Curio et al., 1997; Gobbelé et al., 1998; Halboni et al., 2000; Stenner et al., 2025) were only able to provide isolated brain or spinal cord recordings, thus precluding systematic assessments of CNS-level-specific HFO characteristics within a participant. The robustness and generalizability of our novel approach was ensured by replication analyses on an independent dataset and the assessment of upper as well as lower limb representations, going far beyond previous work.

An important aspect of our study was the application of spatial filtering techniques (based on a variant of canonical correlation analysis, CCA): using such an approach substantially improved the detectability of HFOs across participants (allowing to observe HFOs across all levels in almost every participant after median nerve stimulation) compared to using sensor-level data. This suggests that single-participant HFO studies – e.g. via longitudinal dense-sampling approaches – are feasible in the future with the here-developed methods (which, in contrast to other non-invasive studies (Chander et al., 2022; Stenner et al., 2025), create spatial filters at the individual level). Three further aspects are important in this regard: i) this CCA-based HFO detection was based on a stringent and automated component selection procedure (eliminating experimenter degrees of freedom), ii) it fared worse in the case of tibial nerve responses in the lumbar spinal cord (where only slightly more than half of all participants showed detectable HFOs), and iii) while CCA might be prone to a certain degree of overfitting (Helmer et al., 2024), we quantitatively addressed this issue via validation analyses involving surrogate resting-state data.

We also probed the current limits of non-invasive entire-CNS HFO recordings by assessing the feasibility of detecting HFOs in response to stimulation of sensory nerves with low fiber-count, where a >5-fold reduction of activated nerve-fibres is present when stimulating a finger compared to stimulating the median nerve at the wrist ((Corniani and Saal, 2020); note that other forms of sensory nerve stimulation may give rise to larger responses (Eisen and Elleker, 1980)). Here we noticed that i) cortical HFOs were commonly detectable (see also (Desmedt and Bourguet, 1985; Ozaki and Hashimoto, 2011)) ii) subcortical responses could be assessed when two fingers were stimulated concurrently iii) spinal HFOs could not be assessed at all and iv) detecting HFOs to toe stimulation was not possible at any CNS level. The inability to detect spinal HFOs after finger stimulation contrasts sharply with invasive epidural spinal recordings (Berić et al., 1986) where sensory nerve HFOs are visible and points towards the future need of low-noise recording set-ups in such challenging scenarios. Importantly though, our approach of training CCA on mixed-nerve data (to address the low SNR of the sensory-nerve data) could serve as a template when aiming to investigate HFOs to natural tactile stimulation.

### Characteristics of HFOs across CNS levels

Our approach allowed us to comprehensively characterize HFOs at spinal, subcortical and cortical levels along several features (i.e., latency, topography, frequency, number of wavelet peaks and relation to low-frequency responses). The group-level envelope timecourses demonstrated temporally staggered peaks across spinal, subcortical and cortical levels, occurring at expected timepoints based on the canonical LF-SEPs for both upper and lower limb stimulation across both datasets. Interestingly, at the spinal level the peak latency occurred largely before the LF-SEP, which could indicate a predominantly presynaptic nature. The cortical and subcortical HFO topographies were highly similar to their low-frequency counterparts, as observed previously (Curio et al., 1994; Stephani et al., 2020), even though we observed that high-frequency and low-frequency signals carry non-redundant information (see below) and must thus stem from either at least partly independent signal-generating neuronal ensembles (see also (Gobbelé et al., 2000; Ogawa et al., 2004)) or from distinct processes in different components of pyramidal cells in cortex (Stephani et al., 2024). Our dense electrode montage also allowed for a thorough spatial characterization of spinal HFOs in the cervical and lumbar cord: spinal signals were located centrally in the respective electrode patch, though a lateral aspect – possibly indicating brachial plexus contributions (Curio et al., 1996; Paku et al., 2023) – was observed in the case of cervical responses (we refrain from interpreting lateralization of lumbar responses due to their low SNR).

Time-frequency assessments in both datasets revealed clear peaks of high-frequency (HF) power that – importantly – were mostly well-separated from low-frequency (LF) power in spinal, subcortical and cortical wideband data, indicating genuine oscillations as opposed to filtering artefacts (Bénar et al., 2010). To our knowledge such a clear non-invasive HF-LF distinction has not yet been reported outside of cortex, though invasive spinal recordings (Insola et al., 2008) match our data, also in terms of the spinal peak frequency of around ∼500Hz observed here. Leveraging our unique within-participant design, we could furthermore show a significant increase in HFO frequencies as one ascends the central nervous system from spinal over subcortical to cortical processing stages (apart from the case of lumbar recordings where only few participants were available) – the underlying neurophysiological reason for this hitherto unknown increase within the 400-800Hz frequency band remains to be determined.

By including upper- and lower-limb stimulation conditions, we furthermore observed that at both subcortical and cortical levels, upper-limb HFOs appeared to have higher frequencies across both datasets.

Considering that single peaks in the HFO wavelet have been linked to single peaks in underlying single-unit peristimulus spike histograms (Telenczuk et al., 2011), we next assessed how the number of wavelet peaks evolves across the CNS. In the case of median nerve stimulation, there was a clear and replicable increase across CNS levels in peak counts, that was furthermore independent of the chosen peak threshold criterion (a feature not observed for tibial nerve stimulation). For median nerve stimulation, we observed 2-6 wavelet peaks at the spinal level, which is well in line with invasive recordings (Berić et al., 1986), as are cortical peak counts of 4-8 (Curio et al., 1994; Ozaki et al., 1998); our tibial nerve results are the first such reports to our knowledge. Importantly, our within-participant design allowed us to demonstrate that this increase is of non-random nature as it shows a significant influence of CNS level on peak counts, possibly indicating variations in spiking behavior across these processing stages. Our novel multi-level approach naturally lends itself to future investigations regarding the interplay of HFOs at different levels of the CNS, which is particularly relevant in the context of recent demonstrations that spinal cord HFOs to median nerve stimulation can be modified by top-down factors (Stenner et al., 2025).

Finally, we demonstrated across both cortex and spinal cord that LF and HF signals are at least partly independent. In a first analysis we observed that the covariation between LF and HF signals across participants was strongly reduced when considering the underlying SNR. In a second analysis that leveraged the increased sensitivity achieved by CCA, we investigated within-participant single-trial dynamics and observed that a highly significant difference in HFO strength was not reflected in LF-SEP strength - where timecourses were virtually identical - and vice versa. Together with earlier data (Chander et al., 2022; Stenner et al., 2025; Waterstraat et al., 2021) the consistent effects observed here for both between- and within-individual analyses at different CNS levels strengthen the idea that HF and LF signals contain non-redundant information, thus offering a rich window into human neurophysiology that should be exploited when investigating factors that could modulate processing at these levels (e.g. top-down effects (Gijsen et al., 2021) or effects of pathology (Coppola, 2004; Inoue et al., 2004)).

### Neurobiological origin of HFOs in the CNS

The neurobiological origins of cortical HFO bursts have already been well characterized via several invasive and non-invasive studies in animal models (e.g. rats: Jones et al., 2000; pigs: Ikeda et al., 2002; primates: Baker et al., 2003) and humans (e.g. Cimatti et al., 2006; Curio, 2000; Hashimoto, 2000; Urasaki et al., 2006), with a general consensus that the early part of the cortical HFO burst relates to activity in thalamocortical axon terminals, while the latter part relates to intracortical spiking (though the exact identity of cortical neuron types remains to be determined (Hu et al., 2023; Klostermann et al., 2000)). Interestingly, most group-level cortical HFO envelope timecourses observed here had a two-peaked shape, which might map onto these entities. With regards to subcortical responses, only few studies assessed HFOs in the brainstem (e.g. Halboni et al., 2000; Insola et al., 2010; Restuccia et al., 2004), though the fast spike-rates of dorsal column nuclei would certainly allow for HFO generation (Calvin and Loeser, 1975). Most research has instead focused on thalamically-generated HFOs, which can either arise from thalamocortical projection neurons or local activity and can clearly be dissociated from cortical HFOs (Hanajima et al., 2004; Klostermann et al., 2000; Urasaki et al., 2006). In our study, subcortical HFO latencies were compatible with both thalamic and brainstem generators in the case of median nerve stimulation, but the latency of the response favoured a brainstem generator in the tibial nerve condition.

Compared to supraspinal regions, HFO assessment in the spinal cord (carried out here for both the cervical and lumbar cord for the first time) is in its infancy: to date, there have only been two non-invasive assessments (Chander et al., 2022; Stenner et al., 2025) and only a few invasive studies in humans (e.g. Berić et al., 1986; Chander et al., 2022; Insola et al., 2008; Jeanmonod et al., 1991; Prestor et al., 1997; Urasaki et al., 2006). As for the cellular substrates that give rise to the here-observed spinal cord HFOs, there are several mutually non-exclusive possibilities: i) activity in afferent terminals upon entry to the dorsal horn (Jeanmonod et al., 1991), ii) complex presynaptic travelling waves in ascending tracts with different conduction velocities (Jeanmonod et al., 1991; Urasaki et al., 2006), and iii) activity of spinal interneurons with spike rates exceeding 500 Hz (Seki et al., 2003). Finally, there could be a contribution from peripheral brachial plexus signals to the early part of an HFO wavelet (Berić et al., 1986; Curio et al., 1993, 1991; Urasaki et al., 2006), considering the latency and topography of median nerve spinal HFOs observed here. Although the overall peak latency of the observed spinal HFOs might favour a pre-synaptic spiking origin of these signals, we cannot unequivocally establish such a link considering that the envelope timecourse also showed heightened HFO activity even after the peak of canonical LF-SEPs in the spinal cord. Here, future HFO studies based on protocols investigating integration (Nierula et al., 2024) or stimulus repetition rate effects (Urasaki et al., 2006) could provide critical insights. Finally, it is worth mentioning that while we were successful in identifying sigma-band HFOs in the spinal cord (i.e. in the range of ∼600Hz), distinct higher frequency HFOs in the kappa-band have been identified in cortex non-invasively (Fedele et al., 2012; Waterstraat et al., 2015), and in the spinal cord invasively (Chander et al., 2022; Insola et al., 2008), with invasive data suggesting that the spinal sigma- and kappa-band activity might map onto pre- and postsynaptic spiking, respectively (Insola et al., 2008). In any case, we observed a clear distinction between spinal HFOs and LF-SEPs, strongly suggestive of different information being carried in these two processing channels.

### Limitations and outlook

Several limitations of our approach deserve to be mentioned. First, our investigation of HFOs was restricted by automatically selecting only one CCA component per CNS level. While this allowed for unbiased component selection, it likely does not do justice to the multitude of neuronal processing steps occurring at each level. Future studies could for example subdivide each temporal window to assess HFOs on the rising and falling flank of an LF-SEP peak (Klostermann et al., 1999; Lenoir et al., 2017), which might allow insights into pre- and postsynaptic HFO components at each CNS level by selectively modulating early and late HFOs. Second, our investigation was limited by the low SNR of HFOs, which for example precluded studying responses in the kappa-frequency band (Fedele et al., 2012) or obtaining robust spinal responses to tibial nerve stimulation. Combining our spatial filtering approach with either low-noise-amplifier technology in EEG (Scheer et al., 2011) or novel SQUID systems in MEG (Waterstraat et al., 2021) could be anticipated to help in such and even more challenging scenarios, such as the sensory nerve stimulation conditions investigated here or recording HFOs to natural tactile stimulation. Third, our multi-channel approach is currently not able to provide spatially resolved insights and could thus benefit from combination with fMRI (Ritter et al., 2008), ideally via techniques with enhanced spatial coverage (Finsterbusch et al., 2013), thus allowing for entire CNS fMRI-HFO recordings. Finally, the insights obtained here solely apply to somatosensory HFOs between 400 and 800 Hz. It is thus unclear to what extent other contexts where fast neuronal signaling in different frequency ranges has been observed – e.g. somatosensory 1kHz HFOs (Fedele et al., 2012) or HFOs in epilepsy below 500Hz (Chen et al., 2021) – would benefit from the methodological approaches developed here.

## Conclusion

In the 100 years since the first electroencephalography (EEG) recording was performed (Mushtaq et al., 2024), EEG research largely focused on signals of cortical origin that typically have frequencies below ∼100Hz and are thought to reflect mass post-synaptic activity. Here, using readily available EEG recording equipment, we have critically widened this scope by combining extensions in terms of temporal processing scales (investigating signals up to ∼800Hz) and CNS coverage (extending from cortex to spinal cord). Considering the established link between HFOs and population spikes in cortex, the here-observed HFOs could serve as a complementary non-invasive window on neural population activity at various CNS processing stages, both in health and disease.

## Materials and Methods

### Data acquisition

The data used in this study derive from two experiments, with both datasets publicly available (Dataset 1: https://openneuro.org/datasets/ds004388, Dataset 2: https://openneuro.org/datasets/ds004389). Each dataset includes simultaneously recorded electroencephalography (EEG), electrospinography (ESG), electroneurography (ENG), electrocardiography (ECG) and respiratory data. All participants provided written informed consent prior to participation and the study was approved by the Ethics Committee of the Medical Faculty at the University of Leipzig. For the ESG recordings in both datasets, two patches of 17 electrodes each were placed on a participant’s neck and lower back: one patch was centred on the cervical spinal cord around an electrode placed over the spinous process of the 6th cervical vertebra and another patch was centred on the lumbar spinal cord around an electrode placed over the spinous process of the 1st lumbar vertebra. The electrodes were arranged along the midline of the vertebral column and extended laterally from there, with the ESG reference electrode being located between the cervical and lumbar electrode patches over the spinous process of the 6th thoracic vertebra. The EEG recordings in Dataset 1 were performed using 64 electrodes with standard positions according to the 10-10 system and referenced to the right mastoid; in Dataset 2 EEG was recorded using 39 electrodes, again referenced to the right mastoid. All data was recorded in an acoustically shielded EEG cabin with a sampling rate of 10kHz and an anti-aliasing filter at 2500Hz with a lower cut-off at 0.16Hz. More details on spinal patch construction as well as electrode placement are available in the original publication (Nierula et al., 2024).

Dataset 1 is comprised of 36 healthy right-handed volunteers (18 female) who received electrical mixed nerve stimulation of the median nerve at the left wrist and of the tibial nerve at the left ankle. Electrical stimulation consisted of a 0.2ms square-wave pulse delivered by constant-current stimulators (DS7A, Digitimer Ltd, Hertfordshire, UK; one stimulator for each nerve) via two bipolar stimulation electrodes. Stimulus intensities were adjusted on an individual level to be just above the motor threshold, but not deemed to be painful. Stimulation occurred with an inter-stimulus interval of 763ms and a uniformly distributed jitter of +/-50ms in steps of 1ms. A total of 2000 stimuli were delivered in alternating blocks of 500 stimuli to either the median or tibial nerve. Further resting state recordings were also performed with eyes open before task-based data acquisition.

Dataset 2 is comprised of 24 healthy right-handed volunteers (12 female), who received electrical mixed nerve stimulation of the median nerve at the left wrist and of the tibial nerve at the left ankle (using the identical equipment as in Dataset 1). In addition to this mixed nerve stimulation, sensory nerve stimulation was also performed by stimulating the fingers or toes using ring electrodes. Finger stimulation occurred in three different ways: to the index finger only (finger1), to the middle finger only (finger2), or to both fingers simultaneously (fingers1&2); the same procedure was used for the toes (toe1, toe2, toes1&2). For mixed nerve stimulation, stimulus intensities were adjusted on an individual level to be just above the motor threshold, while for sensory stimulation conditions the stimulation intensity was set to three times the detection threshold, as determined by the method of limits. In both cases the stimulation was deemed to be non-painful. Stimulation occurred with a different frequency compared to Dataset 1 – here we employed an inter-stimulus interval of 257ms and a uniformly distributed jitter of +/-20ms in steps of 1ms. A total of 2000 stimuli were delivered for each mixed nerve and sensory nerve condition. Resting state recordings with eyes open were also performed before task-based data acquisition.

Unless otherwise mentioned, the subsequent preprocessing and analyses are performed on both Dataset 1 and Dataset 2, with the results from Dataset 2 appearing in the supplement.

### ESG and EEG preprocessing

Unless mentioned otherwise, all analyses were performed using Python 3.9 and MNE (https://mne.tools/stable/index.html; version 1.2.0), an open-source toolbox for analysing human neurophysiological data (Gramfort, 2013). In order to remove stimulation artefacts, data were interpolated between −7ms and 7ms relative to stimulus onset for ESG data, and between −1.5ms and 6ms for EEG data using Piecewise Cubic Hermite Interpolating Polynomials (PCHIP). ESG and EEG data were then down sampled to 5kHz, notch filtered around 50Hz with an FIR zero-phase filter and high-pass filtered above 1Hz with a 2nd order Butterworth zero-phase filter. For ESG data, the cardiac artefact was then removed using signal space projection with 6 projectors (Bailey et al., 2024). Bad data segments were annotated if the signal amplitude in the frequency band from 400Hz to 1400Hz was higher than a certain threshold (the mean amplitude across all ESG/EEG channels, plus or minus three times the standard deviation across all ESG/EEG channels); note that a frequency band up to 1400Hz was used as we originally intended to analyse both sigma and kappa band activity (with kappa activity extending to 1200Hz (Fedele et al., 2012)), but were ultimately not able to investigate kappa band activity due to limited SNR in this frequency range. Trials were then marked for exclusion from further analysis based on these annotations. The data was then band pass filtered from 400Hz to 800Hz using a zero-phase 5^th^ order Butterworth filter to isolate sigma band activity (Fedele et al., 2012).

### Canonical correlation analysis

#### Training

CCA was performed to optimise the SNR of the HFO activity using the Modular EEG Toolkit (MEET; https://github.com/neurophysics/meet (Waterstraat et al., 2015)). CCA is used to find the weights *w*_*x*_ and *w*_*y*_ that mutually maximise the correlation

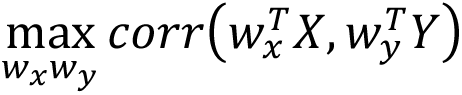

where X and Y are both multi-channel signals. We used a variant of CCA also known as canonical correlation average regression (Waterstraat et al., 2015) – in this case, X is a two-dimensional matrix that contains all single-trial epochs from 1 to N for each channel, and Y contains N times the average SEP for each channel. With this construction, the weight matrix *w*_*x*_ represents the spatial filters that in combination with *w*_*y*_ maximise the correlation between the single trial activity (X) and the average SEP (Y).

The spatial filters *w*_*x*_ were trained using short data segments based on the expected latency and frequency of activity of interest (Table 3), depending on the type of stimulation (median or tibial nerve) and the CNS level under investigation (spinal, subcortical, cortical). For median nerve stimulation, a window of 5ms on either side of the expected peak was used, though in the case of subcortical data, this was reduced to 2.5ms after the expected peak to minimise the effect of cortical activity on the subcortical analysis. For tibial nerve stimulation, due to increased variability of SEP latency across participants, a window of 7.5ms on either side of the expected peak was chosen. Again, in the case of subcortical data, this was reduced to 3.75ms (i.e. 50%) after the expected peak, to remain consistent with the median nerve stimulation analysis.

**Table 3.** Time windows used for CCA training alongside the expected peak latency at each level of the central nervous system.

| **CNS Level** | **Median Nerve Stimulation** |  | **Tibial Nerve Stimulation** |  |
| --- | --- | --- | --- | --- |
|  | Training Window | Expected Peak | Training Window | Expected Peak |
| **Spinal** | 8ms – 18ms | 13ms | 14.5ms – 29.5ms | 22ms |
| **Subcortical** | 9ms – 16.5ms | 14ms | 22.5ms – 33.75ms | 30ms |
| **Cortical** | 15ms – 25ms | 20ms | 32.5ms – 47.5ms | 40ms |

For CCA of spinal data, only electrodes from the relevant spinal patch were included (median nerve: cervical patch; tibial nerve: lumbar patch), while for CCA of subcortical and cortical HFOs every EEG electrode was included. Due to the limited SNR for the sensory nerve conditions in Dataset 2, CCA was trained on data from mixed nerve conditions due to their higher SNR and the spatial filters were then applied to the respective mixed and sensory nerve conditions, similar to previous approaches (Nierula et al., 2024). In each case, the resulting spatial filters were applied to the whole length of the epochs from −100ms to 300ms relative to stimulation.

CCA was also performed on the spinal and cortical data prior to the application of spectral filtering from 400-800Hz, in order to obtain the relevant LF-SEPs alongside the HFOs for mixed nerve conditions. This analysis was not done for subcortical responses due to difficulties in identifying the low frequency somatosensory evoked potential for subcortical responses across participants.

#### Component selection

### High frequency oscillations

In order to prevent possible experimenter-bias in HFO-component selection, we developed an automated approach, whereby a single CCA component was selected from the top four CCA components (ranked by their canonical correlation coefficient) for each participant and each level of the CNS. To select the optimal component, two criteria were considered:

1. the SNR of the HFO must be above 5,
2. the peak of the amplitude envelope of the HFO (as computed with a Hilbert transform) must be within the CCA training time window described in Table 3.

The SNR of the HFO was computed as the absolute value of the peak amplitude within the CCA training window, divided by the standard deviation in a noise period from −100ms to −10ms with respect to stimulation. For sensory nerve stimulation (where responses occur with increased latency (Nierula et al., 2024)), the time window within which the amplitude envelope peak must lie (criterion ii), was the CCA training window shifted rightward by 2ms compared to the mixed nerve condition, and the HFO and its respective envelope was computed across all available conditions combined (i.e. finger1, finger2 and finger12). If there was no component which passed both criteria for a given condition and CNS level, this participant was excluded from further analyses for the given condition and CNS level. If more than one component achieved the specified criteria, the component with the highest SNR was selected.

For each component, the spatial pattern was computed by multiplying the covariance matrix of X by the spatial filters *w*_*x*_, to take the data’s noise structure into account (Haufe et al., 2014). As CCA is insensitive to the polarity of the signal, the optimal component for each participant and condition was manually inspected and marked for inversion if the spatial pattern did not match the expected polarity of a given potential (median: spinal N13, subcortical P14, cortical N20; tibial: spinal N22, subcortical P30, cortical P40).

To demonstrate the benefits of applying spatial filtering, the aforementioned selection criteria were also applied to the data prior to the application of spatial filtering, i.e. selection was performed on data from single channels of interest for median and tibial nerve stimulation respectively (spinal: SC6 and L1, subcortical: CP4 and Cz, cortical: CP4 and Cz after average re-referencing of the data).

### Low frequency somatosensory evoked potentials

In this case, manual selection of the optimal component was performed by selecting the optimal component among the first four components (as ranked by their canonical correlation coefficient) according to the following criteria in accordance with previous research (Nierula et al., 2024; Stephani et al., 2020).

1. The component time course exhibited a peak at the expected latency
2. The component spatial pattern exhibited a topography corresponding to the somatotopy of the stimulated nerve

As CCA is insensitive to the polarity of the signal, if the potential at the expected latency of the time course of the chosen component manifested as a positive peak (spinal N13, spinal N22, cortical N20) or a negative peak (cortical P40), the data was inverted.

#### Validation

We also performed a control analysis to confirm that CCA is identifying genuine stimulus-evoked HFO activity, as CCA will optimise the data in the time window of interest even without genuine task-evoked activity being present. First, *resting-state* EEG and ESG data were pre-processed in the same manner as described previously for task-evoked data, up to and including the isolation of HFOs by spectrally filtering from 400Hz to 800Hz. In order to analyse the resting state data in the same manner as the task-evoked data, the resting state data for each participant was concatenated to itself until the length of the resting state data was equal to, or longer than, that of the task-evoked data. Following this, artificial stimulation triggers were added to the resting state data at timepoints equivalent to their position in the task-evoked data for both the median and tibial nerve condition, before saving these pseudo-task-evoked datasets for each condition. Epochs were then defined spanning from 100ms before to 300ms after the stimulation trigger, with no trials excluded in any participant or condition for either task-evoked or resting state data in this analysis. Following this, during each iteration, 1000 epochs (i.e. 50% of all trials) were subsampled randomly and without replacement from the resting state data and the task-evoked data and submitted to the same CCA procedure as described above for each participant, condition, and CNS level. The first component (as ranked by the canonical correlation coefficient) was selected (without having to satisfy any criteria), and epochs were averaged to form an evoked-response in the case of task-evoked data, and a pseudo evoked-response for resting state data. This process was repeated 1000 times, and after completion, the absolute correlation between each (pseudo) evoked-response and every other (pseudo) evoked-response within the CCA training windows (Table 3) was computed, for resting state and task-evoked data, respectively. The average absolute correlation for each participant, condition and CNS level was then computed by averaging the values in the upper triangle of this 1000×1000 correlation matrix (excluding the main diagonal), to yield a single absolute correlation coefficient in each case. One-sided paired t-tests were then performed to test whether the correlation coefficient was higher in the case of task-evoked data as compared to resting state data within each condition and CNS level.

### Group-level data analyses

#### Timecourses

When presenting group-level timecourses, the amplitude envelopes of single participant HFO bursts were averaged, rather than the timecourses themselves, as slight differences in the HFO burst frequency and latency across participants could lead to cancellation effects – this is prevented by taking the amplitude envelope.

#### Time-frequency representations

Time-frequency representations were obtained for the evoked response for each participant at each CNS level using a Stockwell transform (Stockwell et al., 1996). As CCA was trained using data which was previously band-pass filtered from 400-800Hz, in order to obtain a time-frequency representation across a wider frequency range, the spatial filter of the optimal CCA component was applied to each participants data before band-pass filtering had been performed. This has the added advantage that the data used to generate the time-frequency representations were uncontaminated by potential filter artefacts, such as ringing (Chander et al., 2022).

#### Burst frequency

The burst frequency was quantified for each participant and CNS level (spinal, subcortical, cortical) for which an optimal CCA component was chosen. First, the optimal spatial filter determined by CCA was applied to the broadband electrophysiology data in all channels prior to sigma frequency band isolation, in order to ensure the signal remained uncontaminated by potential filter artefacts. For cortical and subcortical data, channel CP4 was utilised for median nerve stimulation and channel Cz for tibial nerve stimulation. For spinal data, channel SC6 and L1 were used, respectively. These channels represent target locations based on the type of stimulation applied and the processing level under examination. Within these channels of interest, the evoked response was then determined via averaging and subsequently filtered between 400Hz and 800Hz. This filtering was performed to prevent burst frequency detection being affected by the spread of high-power activity from regions outside our interest – this is of particular concern with respect to the low frequency SEP, which has much higher power than the HFOs of interest and can leak into the higher frequency responses. The time-frequency response was determined using a Stockwell transform and these results were then cropped to form a region of interest, defined in the frequency domain from 400Hz to 800Hz, and in the time domain as the CCA training windows (Table 3). This region was then searched for the maximum power, and the frequency associated with this peak was recorded as the burst frequency.

Repeated-measures ANOVAs were performed to determine whether there were differences between the mean burst frequency at different CNS levels in relation to median and tibial nerve stimulation. If differences were detected, post-hoc two-tailed paired t-tests were also performed (using Bonferroni correction for multiple comparisons).

#### Wavelet peak count

The number of wavelets contained within an evoked HFO burst was determined for each participant by counting the number of peaks or troughs which were within 7.5ms on either side of the expected SEP latency (Table 3) for median and tibial nerve stimulation. After obtaining the number of peaks and troughs, these values were added together and divided by 2 to obtain the number of bursts. For the majority of analyses, the time window used for median nerve stimulation is shorter than that for tibial nerve stimulation, due to the decreased temporal variability of responses in relation to median nerve stimulation. For this analysis, however, the same time window was used for both median and tibial nerve stimulation analyses, in order to enable comparison between the wavelet counts in relation to both types of stimulation. Peaks/troughs were counted if they were within the designated time window and had an amplitude of at least X% of the maximum HFO peak/trough amplitude within the same time window, with various thresholds examined (X being 10%, 20%, 25%, 33.33% and 50%).

For each condition and threshold, a non-parametric Friedman test was performed to assess whether there were significant differences in the number of wavelets at each level of the CNS. If significant differences were detected, post-hoc two-tailed Wilcoxon signed-rank tests were performed to identify which CNS levels had significant differences (using Bonferroni correction for multiple comparisons).

#### Sensory nerve stimulation

The ability to detect robust HFOs across participants in response to sensory nerve stimulation at the fingers was tested by means of cluster-based permutation tests for the subcortical and cortical CNS levels. This analysis was not performed for the spinal CNS level after finger stimulation, nor any CNS level after toe stimulation due to the lack of participants with detected HFOs in these cases (all N = 0). For each participant, the amplitude envelope of the HFO timecourse (created by averaging across all available trials for a given participant) was computed by means of a Hilbert transform. In order to allow for performing cluster-based permutation tests of the signal envelope (which is always above zero), the average amplitude envelopes from each participant were individually baseline corrected by calculating the mean of the amplitude envelope in a noise period (−100ms to −10ms relative to stimulation) and subtracting this value from every timepoint in the amplitude envelope. Following this, the corrected individual amplitude envelopes were submitted to a one-sided one-sample cluster-based permutation test, separately for each condition (finger1, finger2, finger12) and CNS level (subcortical, cortical).

#### Relationship between low-frequency and high-frequency responses

Finally, we quantitatively investigated the relationship between the LF-SEP and HFO amplitude in two analyses by asking:

1. across participants, is there a significant correlation between LF-SEP and HFO peak amplitudes (averaging across all trials; resulting in one value for LF-SEP and HFO per participant)
2. within participants, are the same trials exhibiting maximal/minimal responses for LF-SEP and HFO timecourses (considering the top/bottom 10% of trials, as ranked by the single-trial SNR of the HFOs, and separately the SNR of the LF-SEPs)

For the first analysis, within each participant all trials were averaged and the peaks of the evoked low-frequency (LF-SEP) and high-frequency (HFO) responses were extracted for each condition (within the time windows described in Table 3 for both cortical and spinal data). For LF-SEPs, the polarity of the peak was considered (i.e., a negative peak was searched for with respect to the N20, but a positive peak for the P40) as prior to this analysis each timecourse had been inspected after spatial filtering, and if necessary, inverted to match the expected timecourse However, for the HFOs the peak amplitude was obtained irrespective of polarity owing the insensitivity of CCA to polarity. This resulted in a single peak amplitude value for each participant, based on their evoked response computed across all trials for a given condition and CNS level, for both the HFO and LF-SEP. The Pearson correlation between the absolute values of these LF-SEP and HFO peak amplitudes was then computed across participants for each condition and CNS level separately. To address the possibly confounding influence of SNR, we i) computed the SNR for the LF-SEPs and HFOs (as the peak amplitude divided by the standard deviation of a pre-stimulus (−100ms to −10ms) noise period and ii) then calculated partial correlations between the LF-SEP and HFO, when controlling for either the LF-SEP SNR, the HFO SNR or both in tandem. This analysis was not performed for subcortical responses due to difficulties in reliably identifying the LF-SEP peak for subcortical responses across participants.

For the second analysis, within each participant all trials were ranked from highest (top) to lowest (bottom) according to the HFO-SNR for the cortical and spinal data after median and tibial nerve stimulation. Separate evoked timecourses were then created for the LF-SEP and HFO amplitude envelope by averaging across either the top or the bottom 10% of trials. The evoked timecourse for the bottom 10% of trials was then subtracted from the timecourse for the top 10% of trials to form a difference time course for each participant, which were then submitted for statistical assessment at the group level (cluster-based permutation test). This analysis was then repeated based on the LF-SEP SNR rather than the HFO-SNR – in this case, prior to the computation of the LF-SEP SNR each trial was baseline corrected by subtracting the mean value of the data in the entire trial window from −100ms to 299ms from each datapoint in the same trial, excluding the interpolation time period relative to stimulation (−7ms and 7ms relative to stimulus onset for ESG data, and between −1.5ms and 6ms for EEG data). This was done to account for baseline shifts between single trials, prior to ranking.

## Supporting information

Supplementary Results

## Acknowledgements

We would like to thank all volunteers who participated in this study. Additionally, we want to thank everyone who assisted in data acquisition.

## Author Contributions

Listed alphabetically according to CRediT taxonomy (https://credit.niso.org).

Conceptualization: EB, FE, BN, VN, TS

Data curation: BN

Formal analysis: EB

Funding acquisition: FE

Investigation: EB, BN

Methodology: EB, FE, VN

Project administration: EB, FE

Resources: FE, VN

Software: EB

Supervision: FE, VN

Visualization: EB

Writing – original draft: EB, FE

Writing – review & editing: EB, FE, BN, TS, VN, GC, GW

## Data and code availability

The underlying data and code are openly available via OpenNeuro (Dataset 1: https://openneuro.org/datasets/ds004388 , Dataset 2: https://openneuro.org/datasets/ds004389) and GitHub (https://github.com/eippertlab/hfo-cns), respectively.

## Ethics

All participants gave written informed consent. The study was approved by the Ethics Committee of the Medical Faculty of the University of Leipzig.

