## Supplementary Results for "Somatosensory high frequency oscillations across the human central nervous system"

**This file includes:**

Figures S1 to S9
Table S1

### Results

#### Single participant analyses

Similar to Dataset 1, we also observed distinct HFO bursts at each CNS level in Dataset 2, as shown in representative individual timecourses (Figure S1).

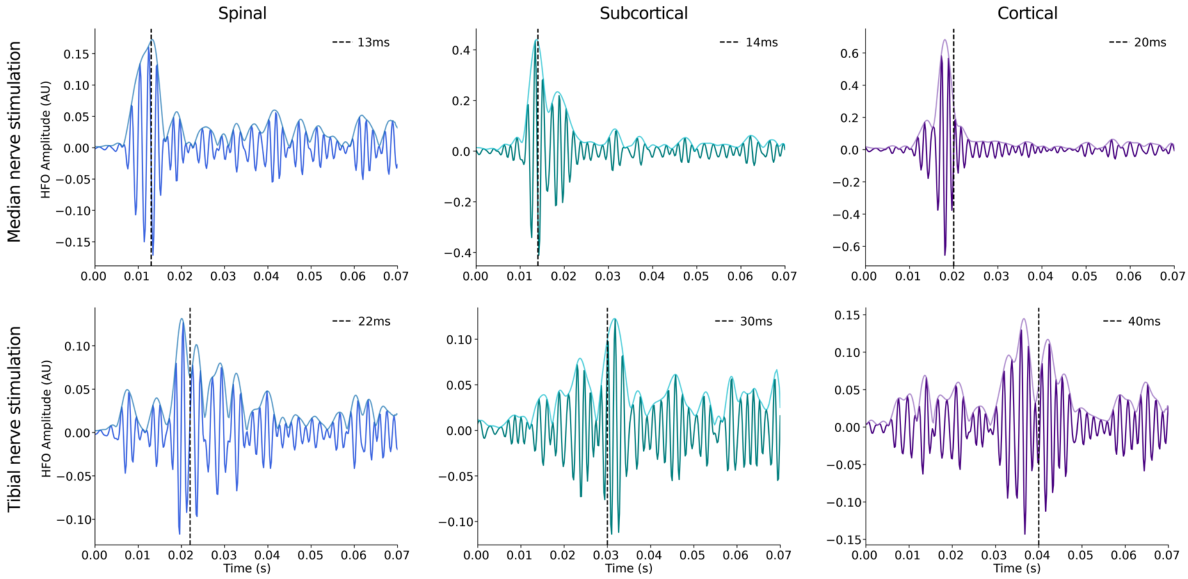


**Figure S1. Individual HFO bursts across the CNS.** Single participant HFO timecourses (thick lines) and amplitude envelopes (thin lines) in response to median nerve stimulation (top) and tibial nerve stimulation (bottom) across spinal (left), subcortical (middle) and cortical (right) recordings (participant 5). For each panel, the canonical latency of the LF-SEP is displayed as a dashed vertical line.

#### Group level timecourses and topographies

We observed similar group-level HFO envelope timecourses for both median and tibial nerve stimulation in Dataset 2 (Figure S2) as in Dataset 1. The group-level timecourses showed peaks at latencies that are in accordance with the CNS level, with peak latencies for median nerve stimulation at 11.2ms (spinal), 14.0ms (subcortical) and 18.0ms (cortical), and peak latencies for tibial nerve stimulation at 22.6ms (spinal), 28.6ms (subcortical) and 37.4ms (cortical).

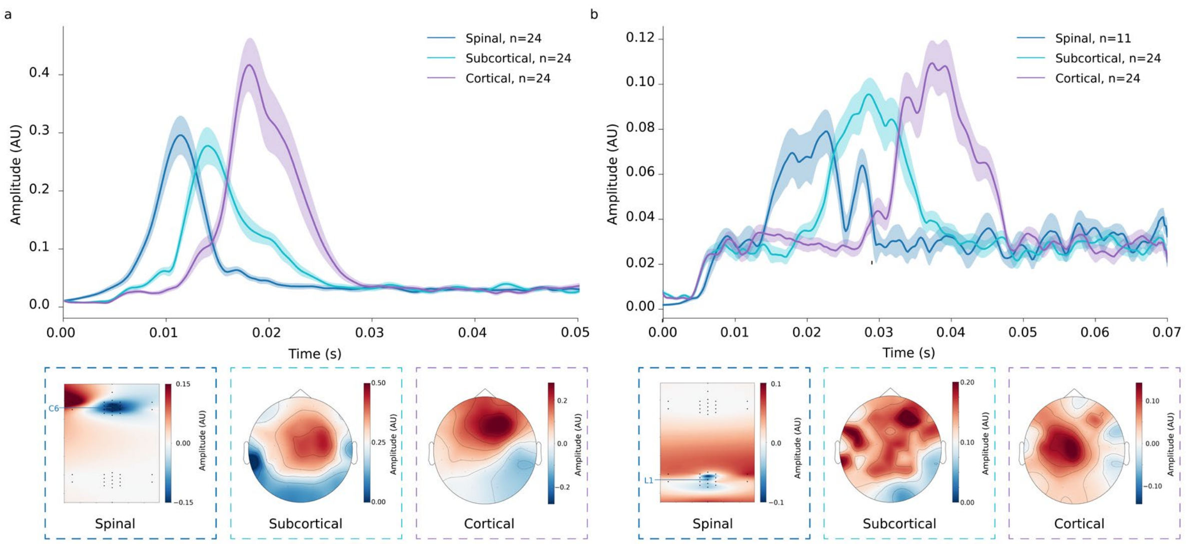


**Figure S2. Group-level HFO timecourses and topographies across the CNS.** Depicted are group-averaged HFO amplitude envelopes and the associated spatial topographies for spinal, subcortical and cortical responses to a) median nerve stimulation and b) tibial nerve stimulation. The error bands reflect the SEM across participants and the number of eligible participants entering each average is given in the legend. Please also note the different scales applied to each plot.

### Cortical and subcortical timecourse in sensor space across all participants

To support the CCA-based detection of HFOs at latencies suggestive of subcortical origin, we also provide evidence for their existence in single-channel sensor space data (Figure S3).

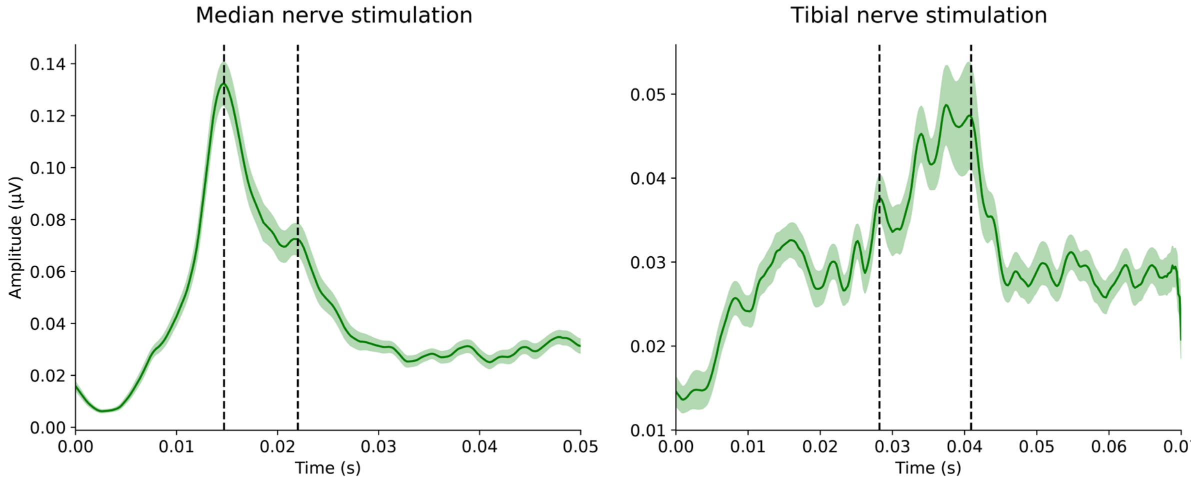


**Figure S3**. **Group-level sensor space timecourses.** HFO amplitude envelopes, after median nerve stimulation (channel CP4) and tibial nerve stimulation (channel Cz) across both datasets (n=60); no spatial filtering via CCA has taken place. In both cases, the HFO amplitude envelopes for each individual participant has been aligned based on the timing of the cortical low frequency SEP (median: N20, tibial: P40). For median nerve stimulation, peaks can be seen at 14.7ms and 22.0ms (dashed lines) – timings typical for subcortical and cortical activity, respectively. For tibial nerve stimulation, similar peaks related to subcortical and cortical activity can be seen at 28.2ms and 40.9ms (dashed lines), respectively (though with a much higher noise-level).

#### Low frequency and high frequency spatial topographies

In order to demonstrate the similarity of low frequency and high frequency spatial topographies at all CNS levels (Dataset 1), we depict both of these here (Figure S4).

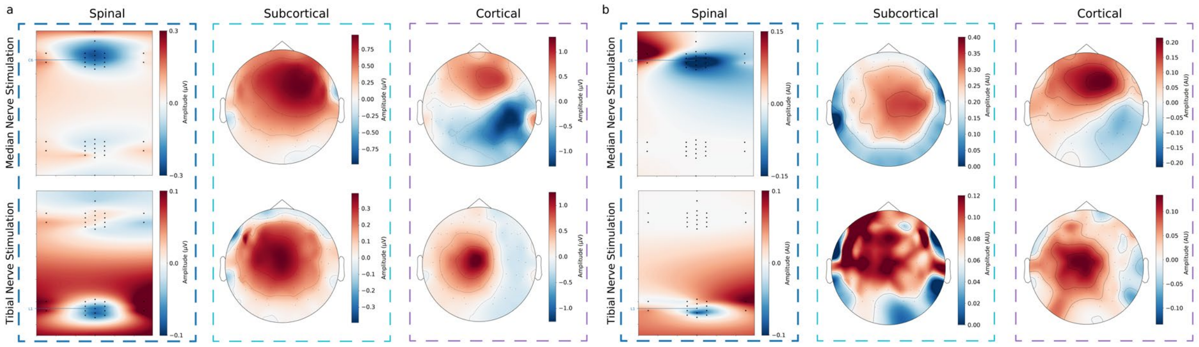


**Figure S4. Group level spatial topographies for low frequency and high frequency responses**. For low frequency spatial topographies (a) the spinal responses are referenced to electrode TH6 (at the spinous process of the sixth thoracic vertebra), subcortical responses are referenced to electrode RM (right mastoid), and cortical responses have been average re-referenced. For high frequency spatial topographies (b), the images shown represent the average across the spatial patterns associated with the chosen CCA component for each participant. Note that in the case of spinal responses, only the relevant electrode patch (either cervical or lumbar spinal cord) was submitted to CCA.

#### CCA validation analyses

Here, we provide detailed statistical results of the CCA validation analyses carried out on task-evoked and resting state data for both Dataset 1 and Dataset 2. Out of the 12 performed analyses (2 datasets x 2 stimulation conditions x 3 CNS levels), 10 analyses showed significantly stronger effects in the stimulation-based data (Table S1); only subcortical and spinal HFOs after tibial nerve stimulation in Dataset 2 failed to pass the significance threshold, possibly also due to power issues caused by the lower participant numbers (N = 11 for spinal level).

**Table S1. CCA validation results.** Provided are the mean ± standard error of the mean correlation across participants for each CNS level (cortical, subcortical, spinal) and condition (median nerve, tibial nerve) for both task-evoked and resting state data in Datasets 1 and 2. Further, for each CNS level and condition, the p-value resulting from each paired t-test is indicated, showing in which instances the correlation for task-evoked data is significantly greater than the resting state data.

| | | Dataset 1 | | | Dataset 2 | | |
|---|---|---|---|---|---|---|---|
| | | Task-Evoked | Resting State | P-value | Task Evoked | Resting State | P-value |
| **Cortical** | Median Nerve | 0.95 ± 0.02 | 0.52 ± 0.01 | <0.001 | 0.93 ± 0.03 | 0.49 ± 0.02 | <0.001 |
| | Tibial Nerve | 0.58 ± 0.03 | 0.46 ± 0.01 | <0.001 | 0.52 ± 0.03 | 0.42 ± 0.01 | <0.05 |
| **Subcortical** | Median Nerve | 0.89 ± 0.02 | 0.57 ± 0.01 | <0.001 | 0.90 ± 0.02 | 0.54 ± 0.02 | <0.001 |
| | Tibial Nerve | 0.54 ± 0.02 | 0.49 ± 0.01 | <0.05 | 0.48 ± 0.02 | 0.45 ± 0.01 | 0.14 |
| **Spinal** | Median Nerve | 0.91 ± 0.02 | 0.46 ± 0.01 | <0.001 | 0.88 ± 0.03 | 0.48 ± 0.01 | <0.001 |
| | Tibial Nerve | 0.43 ± 0.01 | 0.40 ± 0.01 | <0.05 | 0.39 ± 0.01 | 0.41 ± 0.01 | 0.91 |

#### Group level time-frequency representation

A group-level spectral characterization of HFOs in broadband data from Dataset 2 revealed results highly similar to Dataset 1, with a clear spectral separation of high frequency and low frequency peaks at all CNS levels (Figure S5).

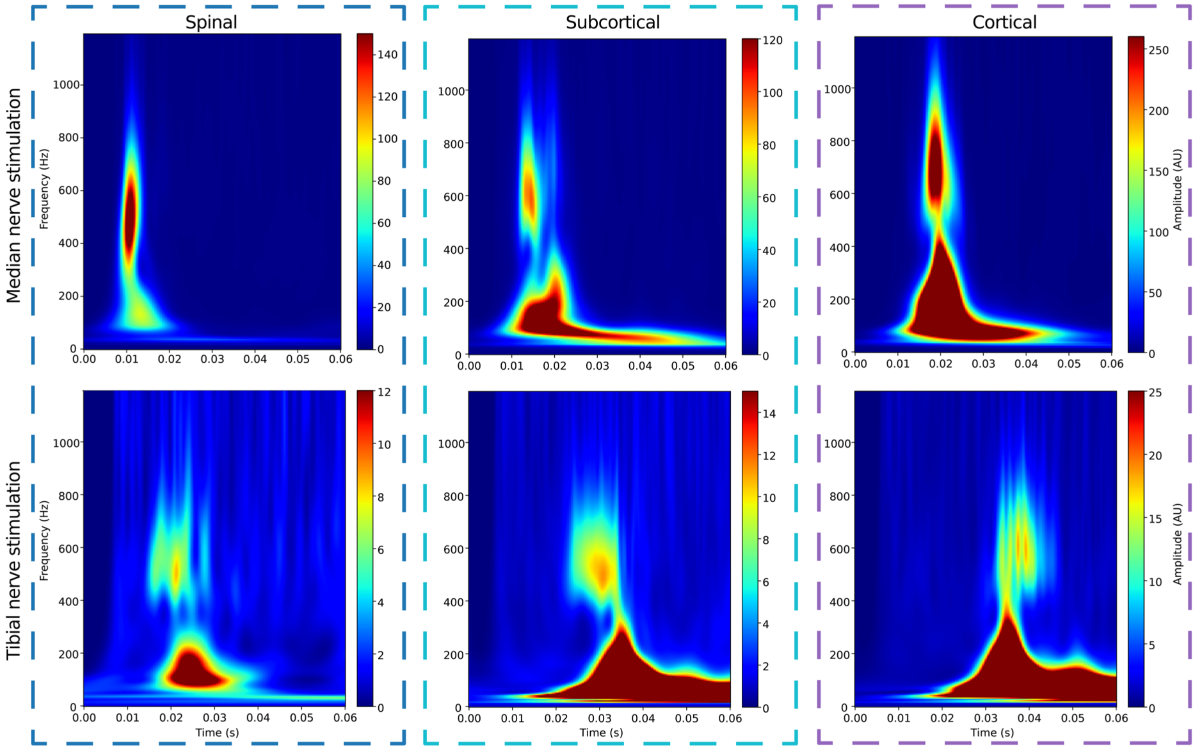


**Figure S5. Evoked power across the CNS.** Group-average time-frequency representations of the evoked response to median nerve (top) and tibial nerve (bottom) stimulation, including spinal (left), subcortical (middle) and cortical (right) responses. Please note the different scales applied to each time-frequency plot.

#### Burst frequency

Statistical testing via repeated measures ANOVAs revealed significant differences between the mean burst frequency at different CNS levels in response to median nerve stimulation in both datasets. Follow-up post-hoc paired t-tests were thus performed and revealed that for participants that have an optimal CCA component identified at all levels of the CNS (N=33 in Dataset 1, n=24 in Dataset 2), the spinal burst frequency is significantly lower than the burst frequency at the subcortical level ($t=-7.54$ for Dataset 1 and $t= -6.49$ for Dataset 2, both $p<0.05$), and the burst frequency at the subcortical level is significantly lower than the cortical burst frequency ($t=-8.70$ for Dataset 1 and $t=-8.23$ for Dataset 2, both $p<0.05$).

In response to tibial nerve stimulation, a similar repeated-measures ANOVA failed to reach significance ($p=0.055$ for Dataset 1 and $p=0.09$ for Dataset 2), possibly influenced by a lack of power due to the lower number of participants having an optimal CCA component at each level.

#### Wavelet peak count

##### Simulation analysis to investigate filter artefacts

The demonstration that spinal, subcortical and cortical HFOs can be observed in broadband data (Dataset 1: Figure 3; Dataset 2: Figure S5) rules out the possibility that HFOs are due to ringing artefacts induced by band-pass filtering. However, to assess whether – in the band-pass filtered data – specific HFO characteristics, such as the reported number of wavelet peaks might be affected by filter ringing to a certain degree, simulations were carried out with a synthetic signal. This signal consisted of a unit impulse summed with 1/f noise. The standard deviation of the simulated noise was set to match that of the noise in the baseline period in either the empirical data prior to band-pass filtering from 400Hz to 800Hz or in the empirical data after band-pass filtering from 400Hz to 800Hz – the latter serving as a typical SNR which is measured in the frequency band of interest. Such separate simulations were necessary as it is not feasible to simultaneously satisfy both noise criteria with a single simulation. This is likely due to additional noise sources present in the empirical data, such as physiological noise from cardiac activity (Bailey et al., 2024) and muscle activity (Ma et al., 2012), which do not adhere to the strict 1/f pattern.

We first consider how to estimate the noise level for the simulation that matches data before band-pass filtering from 400Hz to 800Hz. In the empirical data – averaged across both conditions and both datasets – we observed the group-average peak amplitude of the low-frequency somatosensory evoked potential to be 0.4μV in spinal data, and 2.2μV in cortical data (note we use the empirical data's evoked response averaged across 2000 trials to inform the simulations, as the single trial signal amplitudes cannot be robustly estimated from the empirical single-channel data). The standard deviation in the baseline period (-100ms to -10ms relative to stimulation) prior to averaging across all trials was observed to be 5.6μV in spinal data, and 11.3μV in cortical data. Since we aimed to have the signal-to-noise ratio (SNR, i.e. signal amplitude divided by pre-stimulus noise standard deviation) of the simulated signal (unit impulse + noise, signal length of 2000 samples) to match the SNR in our empirical data we needed to find the scaling factor x, that would allow us to match the SNR of each single trial in our simulated signal (1/x) to that of our spinal and cortical empirical data. Thus, since

Spinal:

$$\frac{1}{x} = \frac{0.4\mu V}{5.6\mu V}$$
$$x = 14.0$$

Cortical:

$$\frac{1}{x} = \frac{2.2\mu V}{11.3\mu V}$$
$$x = 5.1$$

the 1/f noise (with a standard deviation in the baseline period of 1 prior to scaling) was scaled up by a scaling factor 14.0 (to match spinal data) or 5.1 (to match cortical data) before being added to the unit impulse to form the synthetic signal.

Next, we considered how to estimate the scaling factor such that the noise level for the simulation after bandpass filtering from 400-800Hz matches that of the empirical data after band-pass filtering from 400Hz to 800Hz. Since we observed the standard deviation of the noise in the band-pass filtered empirical data for both cortical and spinal data to be 3.5μV, we scaled the 1/f noise by 12.0. In this case, we only matched the standard deviation of the noise, rather than employing the SNR calculation shown above, as the degree to which the signal we encounter after band-pass filtering is introduced by filter ringing is the object of this study, and thus not reliable to inform the degree to which the noise should be scaled.

The process of generating 1/f noise and scaling was repeated 2000 times for each scaling factor, to simulate 2000 trials per participant. Following this, the synthetic signals were band-pass filtered from 400Hz to 800Hz using the same filter as in the main analysis (zero-phase $5^{th}$ order Butterworth filter, in forward and backward directions). The resulting synthetic signals after filtering were averaged across trials and the SNR of the simulated data was determined by finding the peak magnitude within 50 samples on either side of the unit impulse location (to match the number of samples included for the equivalent peak search in the main study) and dividing this by the standard deviation in the baseline period (from sample 10 to sample 461 to match the number of samples used in the equivalent calculation in the main study). The computed SNR in the 400-800Hz frequency range was then compared to the SNR threshold of 5 (also in the 400-800Hz frequency range), as this was the criterion we required in the main empirical analysis to retain a participant’s data for further analysis (see section 4.3.2).

All simulations were performed a total of 60 times (to match our empirical participant numbers; Dataset 1: N=36, Dataset 2: N=24). In none of the simulations did the SNR exceed the threshold of 5: the SNR of the simulated evoked response after filtering from 400-800Hz across 60 iterations was 2.00±0.67 (mean ± standard deviation) for scaling factor 14.0 (matching low-frequency spinal empirical data), 2.90±0.84 for scaling factor 5.1 (matching low-frequency cortical empirical data), and 2.14±0.68 for scaling factor 12.0 (matching high-pass filtered spinal and cortical empirical data). Comparing this to the empirical results reported in the main analysis (Table 1) establishes that characteristics of the HFOs we observe in the main analysis are unlikely to be due to filter ringing. This is likely due to our signal in the raw electrophysiological data either not being sufficiently sharp or not having a large-enough SNR to cause significant filter ringing above the noise floor, an idea supported by previous work (Widmann and Schröger, 2012).

##### Statistical analysis of differences across CNS levels

The same qualitative pattern of results was observed for both Dataset 1 in Figure 4, and Dataset 2 in Figure S6 – i.e. an increase in the number of peaks from spinal over subcortical to cortical levels for median nerve stimulation, with no clear trend for responses to tibial nerve stimulation.

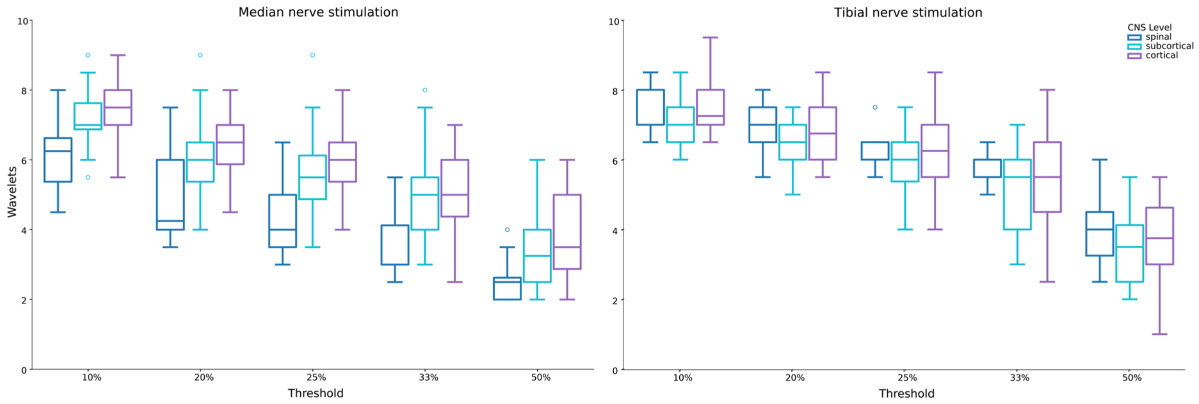


**Figure S6. Wavelet peak count across the CNS.** Depicted are the group-average number of wavelet peak bursts at each level of the CNS, dependent on different thresholds used for peak-counting (i.e. relation of peak/trough size to maximum peak/trough, respectively).

This qualitative observation was tested formally via non-parametric Friedman tests for each Dataset. In Dataset 1, for both median and tibial nerve stimulation there was a significant effect of CNS-level on the number of peaks at all threshold levels. Follow-up post-hoc Wilcoxon tests further revealed significant differences between spinal-subcortical, and subcortical-cortical wavelet peak counts at each threshold level for median nerve stimulation (all $p<0.05$ after Bonferroni correction), but no significant effects for tibial nerve stimulation. For Dataset 2, there was again a significant effect of CNS-level on the number of peaks at all threshold levels for both median and tibial nerve stimulation. Follow-up post-hoc Wilcoxon tests further revealed significant differences between spinal-subcortical, and subcortical-cortical wavelet peak counts at each threshold level for median nerve stimulation (all $p<0.05$ after Bonferroni correction), but no significant effects for tibial nerve stimulation.

Importantly, these results held across the various thresholds for peak definition we examined (10%, 20%, 25%, 33.33% and 50%), pointing to a robust effect that is not dependent on peak-detection criteria (additional analyses further did not provide evidence for a relation between the signal-to-noise ratio of the HFO response and the wavelet peak count).

#### Sensory nerve stimulation

Cluster-based permutation testing revealed that robust group-level cortical responses can be detected after single (finger1, finger2) and double digit (finger12) stimulation, as well as subcortical responses after double digit stimulation. For cortical responses, a significant cluster was identified for finger1 stimulation from 21ms to 33ms ($p=0.006$), for finger2 stimulation from 18ms to 31ms ($p=0.0015$) and for finger12 stimulation from 15ms to 33ms ($p=0.0005$). For subcortical responses, a significant cluster was identified for finger12 stimulation from 15ms to 25ms ($p=0.016$), while strong but non-significant clusters are identified for finger1 ($p=0.063$) and finger2 ($p=0.086$) conditions from 15-19ms and 15-18ms, respectively.

#### Relationship between low-frequency and high-frequency responses

##### Correlation between average LF-SEP and HFO peak amplitude

In Dataset 1, significant LF-HF correlations were observed for median-nerve stimulation evoked spinal ($r_{N13} = 0.66$, $p_{N13} < 0.001$) and cortical responses ($r_{N20} = 0.38$, $p_{N20} < 0.05$), but not for tibial-nerve stimulation evoked responses ($r_{N22} = 0.30$, $p_{N22} = 0.19$; $r_{P40} = 0.19$, $p_{P40} = 0.26$). Importantly, when accounting for the possibly confounding effects of SNR (by considering the partial correlation controlling for LF-SEP SNR, HFO SNR or both), only the spinal, tibial condition controlling for both HFO and LF-SEP SNR showed a significant result ($r_{partial,\ N22} = 0.54$, $p<0.05$).

In Dataset 2, significant LF-HF correlations were observed for median-nerve stimulation evoked spinal ($r_{N13} = 0.67$, $p_{N13} < 0.001$) and cortical responses ($r_{N20} = 0.23$, $p_{N20} = 0.29$), but not for tibial-nerve stimulation evoked responses ($r_{N22} = -0.40$, $p_{N22} = 0.23$; $r_{P40} = 0.26$, $p_{P40} = 0.23$). Additionally, none of these correlations were significant at the $p < 0.05$ level when controlling for the confounding effects of SNR (LF-SEP SNR, HFO SNR or both).

##### Comparing LF-SEP and HFO timecourses when considering the strongest versus the weakest trials

For the HFO-SNR based ranking in Dataset 1, in spinal data, there was a significant difference in the amplitude envelopes formed using the strongest versus the weakest trials for the HFO data between 7ms and 19ms for median nerve stimulation, and between 14ms and 29ms for tibial nerve stimulation (both $p=0.001$), while there was no significant difference in the relevant low frequency SEP traces (not even non-significant clusters were detected). A similar effect was found for cortical data, (significant cluster between 13ms and 26ms for median nerve stimulation and between

32ms and 49ms for tibial nerve stimulation; both p=0.001), while there was again no statistically significant difference in the relevant low frequency SEP traces. When considering trials ranked according to the LF-SEP SNR in Dataset 1, statistically significant differences were observed between the strongest and weakest trials in terms of their LF-SEPs, but not their HFOs (Figure S7). For spinal data, there was a significant difference in the LF-SEP data between 7ms and 31ms for median nerve stimulation, and between 7ms and 45ms for tibial nerve stimulation (both $p < 0.02$), while there was no significant difference in the relevant HFO amplitude envelopes. A similar effect was found for cortical data, (significant cluster between 6ms and 39ms for median nerve stimulation and between 6ms and 63ms for tibial nerve stimulation; both $p = 0.001$).

The pattern of results observed in Dataset 1 was replicated in Dataset 2. First, for the HFO SNR based ranking, cluster-based permutation tests revealed that for spinal data, there was a significant difference in the amplitude envelopes formed using the strongest or the weakest trials for the HFO data between 7ms and 22ms for median nerve stimulation, and between 15ms and 24ms, as well as 25ms and 30ms, for tibial nerve stimulation (all $p < 0.004$), while there was no significant difference in the relevant low frequency SEP traces (not even non-significant clusters were detected). A similar effect was found for cortical data, (significant cluster between 12ms and 27ms for median nerve stimulation and between 32ms and 48ms for tibial nerve stimulation; both $p = 0.001$), while there were again no statistically significant differences in the relevant low frequency SEP traces. Second, for spinal data, there was a significant difference in the LF-SEP data between 7ms and 22ms for median nerve stimulation, and between 7ms and 49ms for tibial nerve stimulation (both $p < 0.025$), while there were no significant differences in the relevant HFO amplitude envelopes. A similar effect was found for cortical data, (significant cluster between 6ms and 36ms for median nerve stimulation and between 17ms and 68ms for tibial nerve stimulation; both $p < 0.01$).

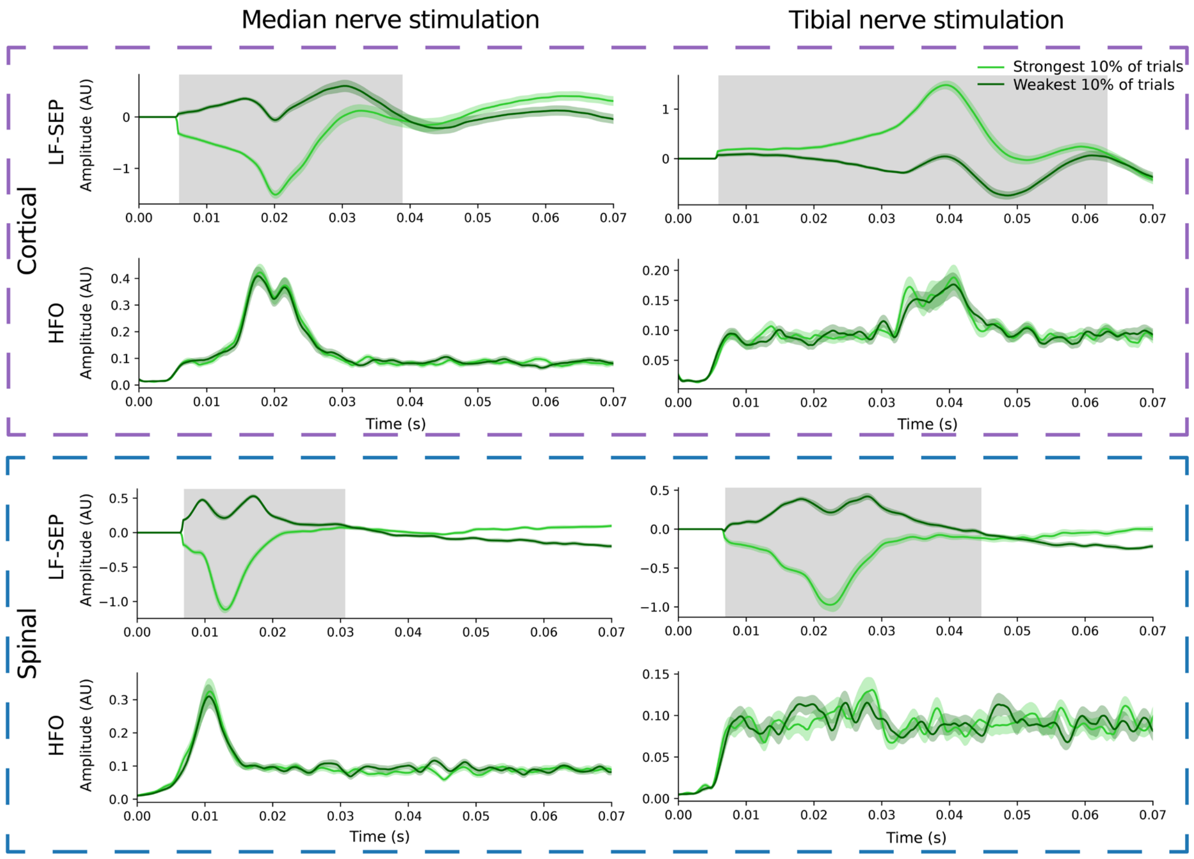


**Figure S7. Difference between low-frequency and high-frequency signals as ranked according to the LF-SEP SNR (Dataset 1).** Depicted are the group-average low-frequency somatosensory evoked potentials (LF-SEP) and group-average high-frequency amplitude envelopes (HFO) when considering either the strongest (light green) or weakest (dark green) 10% of trials. The upper two panels show cortical data after median (left; n=36) and tibial (right; n=36) nerve stimulation, whereas the lower two panels show spinal data (median nerve: n=35; tibial nerve: n=20). The grey areas indicate significant clusters at $p < 0.05$ from a cluster-based permutation test. Please note the different y-axis scales employed in each plot.

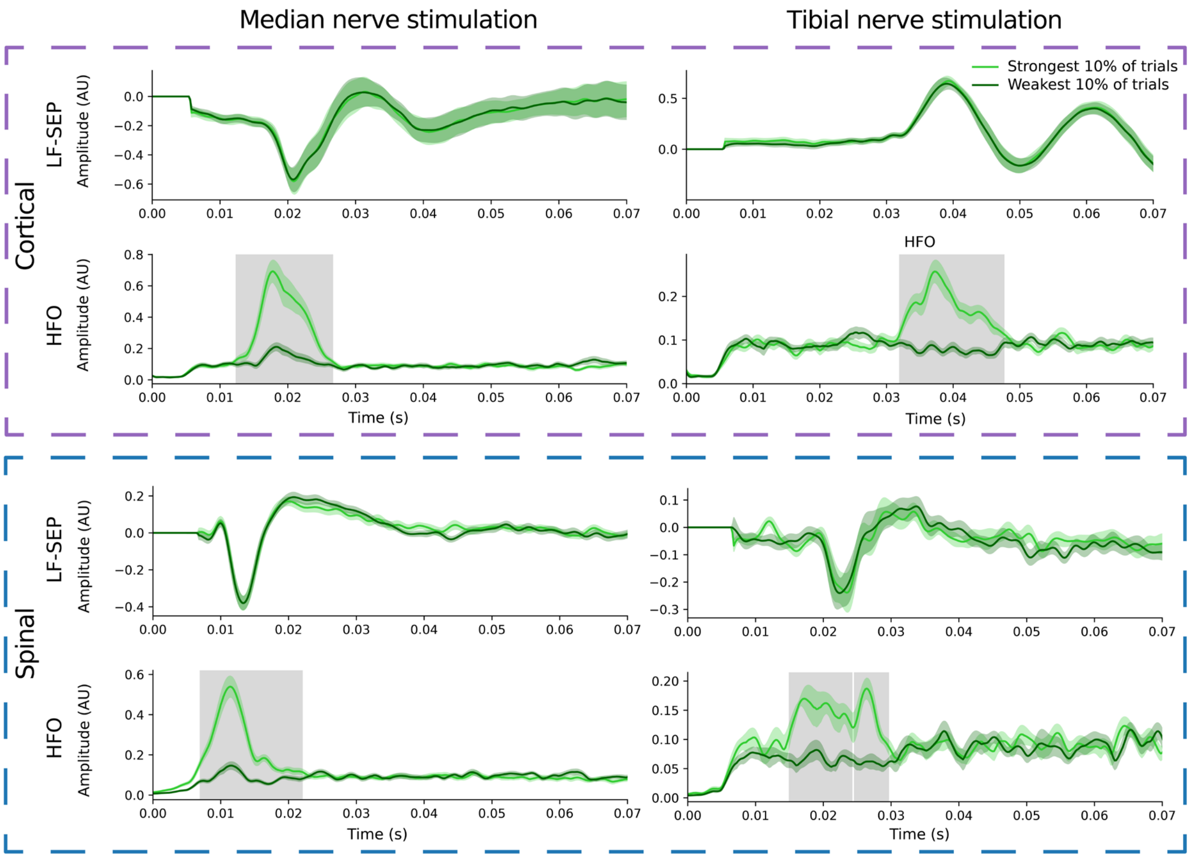


**Figure S8. Difference between low-frequency and high-frequency signals as ranked according to the HFO SNR (Dataset 2).** Depicted are the group-average low-frequency somatosensory evoked potentials (LF-SEP) and group-average high-frequency amplitude envelopes (HFO) when considering either the strongest (light green) or weakest (dark green) 10% of trials. The upper two panels show cortical data after median (left; n=23) and tibial (right; n=23) nerve stimulation, whereas the lower two panels show spinal data (median nerve: n=24; tibial nerve: n=11). The grey areas indicate significant clusters at $p < 0.05$ from a cluster-based permutation test. Please note the different y-axis scales employed in each plot.

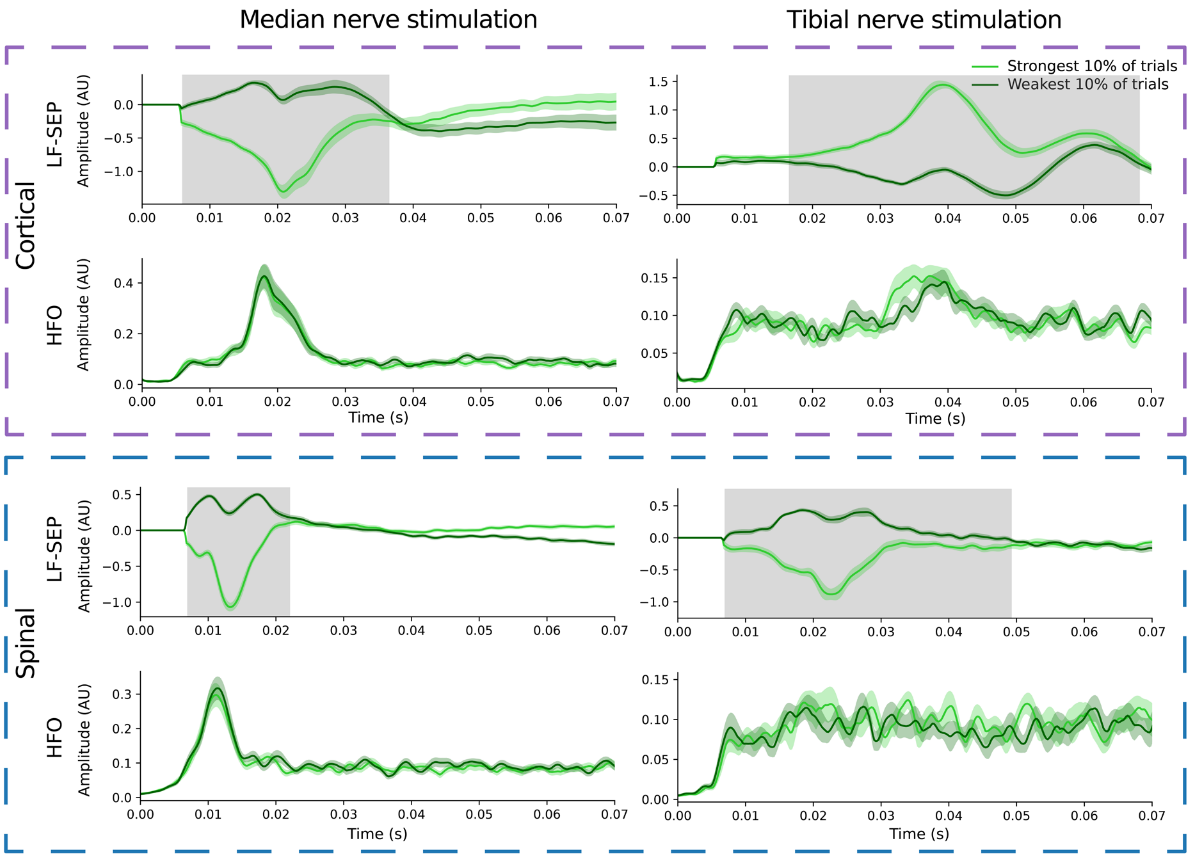


**Figure S9. Difference between low-frequency and high-frequency signals as ranked according to the LF-SEP SNR (Dataset 2).** Depicted are the group-average low-frequency somatosensory evoked potentials (LF-SEP) and group-average high-frequency amplitude envelopes (HFO) when considering either the strongest (light green) or weakest (dark green) 10% of trials. The upper two panels show cortical data after median (left; n=23) and tibial (right; n=23) nerve stimulation, whereas the lower two panels show spinal data (median nerve: n=24; tibial nerve: n=11). The grey areas indicate significant clusters at $p < 0.05$ from a cluster-based permutation test. Please note the different y-axis scales employed in each plot.
